# A Novel and Functionally Diverse Class of Acetylcholine-gated Ion Channels

**DOI:** 10.1101/2021.10.06.463318

**Authors:** Iris Hardege, Julia Morud, Amy Courtney, William R Schafer

## Abstract

Fast cholinergic neurotransmission is mediated by acetylcholine-gated ion channels; in particular, excitatory nicotinic acetylcholine receptors play well established roles in virtually all nervous systems. Acetylcholine-gated inhibitory channels have also been identified in some invertebrate phyla, yet their roles in the nervous system are less well understood. We report the existence of multiple new inhibitory acetylcholine-gated ion channels with diverse ligand binding properties in *C. elegans*. We identify three channels, LGC-40, LGC-57 and LGC-58, whose primary ligand is choline rather than acetylcholine, as well as the first evidence of a truly polymodal channel, LGC-39, which is activated by both cholinergic and aminergic ligands. Using our newly deorphanised channel evidence we uncover the surprising extent to which a single neuron expresses both excitatory and inhibitory channels, not only for acetylcholine but also the other major neurotransmitters. The results presented in this study offer a new insight into the potential evolutionary benefit of a vast and diverse repertoire of LGICs to generate complexity in an anatomically compact nervous system.

## Introduction

Rapid signalling through neuronal networks is essential for producing coordinated behaviours in animals. At the fundamental level, fast neuronal transmission is mediated through neurotransmitter release resulting in activation of ion channels on the postsynaptic neuron. In the textbook view, based predominately on mammalian systems, there are two major excitatory neurotransmitters, glutamate, and acetylcholine, and two inhibitory neurotransmitters, GABA and glycine. Glutamate acts through a family of tetrameric ligand-gated cation channels, while the remaining neurotransmitters activate pentameric receptors from the Cys-loop receptor superfamily of ligand-gated ion channels (LGICs). Although LGICs are highly conserved across phyla, ligand binding properties and ion selectivity diverge significantly, resulting in a large diversity of mechanisms by which small molecules acting via LGICs can influence the activity in neuronal circuits.

Despite their conserved general structure, Cys-loop LGICs vary significantly in their functional properties, particularly when channels from invertebrate phyla are considered. For example, insects and nematodes express inhibitory glutamate receptors from the Cys-loop superfamily, localised both in neurons and muscles, which are targets of the anthelminthic drug ivermectin (Cully et al., 1996, 1994). Many animals including insects, nematodes and mammals also express LGICs which can be gated by aminergic ligands, including histamine-gated chloride channels (Gisselmann et al., 2002), important for fly visual processing, a number of nematode channels, involved in learning and motor control (Morud et al., 2021; Pirri et al., 2009), as well as the excitatory mammalian 5-HT_3_ receptor (Kondo et al., 2014; Lombaert et al., 2018). Even more divergent roles for LGICs have been identified in marine species, where LGICs gated by terpenes and chloroquine function as chemoreceptors in octopus (van Giesen et al., 2020).

Cys-loop channels can be subdivided into two large clades, the first containing the nicotinic receptors and their paralogues including the 5-HT_3_ receptors, and the other containing channels more closely related to GABA_A_ receptors (Jones and Sattelle, 2008). The *C. elegans* genome contains several subfamilies which appear to have diversified independently from vertebrate channels during evolution, leading to the existence of several nematode specific LGIC subfamilies. The *C. elegans* genome contains a number of GABA_A_-like subfamilies, including genes encoding both anion and cation-selective channels (Jobson et al., 2015; Margie et al., 2013; Putrenko et al., 2005; Ranganathan et al., 2000; Ringstad et al., 2009; Yassin et al., 2001). One of these subfamilies consist of genes for both anion and cation selective monoamine-gated channels, another of acetylcholine-gated anion channels or channels that are still largely uncharacterised. One member in one of these subgroups, LGC-40, has previously been reported to be a low affinity serotonin-gated channel also gated by acetylcholine and choline (Ringstad et al., 2009). The properties of the remaining channels in these subgroups, including their ligands, ion selectivity, and expression patterns, are currently unknown.

Here we describe the deorphanisation and characterisation of five new *C. elegans* LGICs activated by acetylcholine. One of these, LGC-39, forms a polymodal homomeric anion channel activated by acetylcholine as well as by monoamines, while the others are activated by choline and/or acetylcholine. Using public single cell RNAseq expression data together with our new electrophysiological data, we also predict the polarity of synapses in the worm connectome, as well as intracellular localisation patterns for uncharacterised LGICs. These results highlight the unexpected functional diversity of cholinergic signalling in the *C. elegans* nervous system.

## Results

### Deorphanisation of Uncharacterised LGICs Reveals Diversity of Cholinergic Channels

The *C. elegans* genome encodes a diverse superfamily of pentameric ligand-gated ion channels (LGICs), of which several subfamilies are poorly characterised. Here we investigate the diverse group, which consists of 3 subgroups named after a channel from each group; the LGC-45 group, the LGC-41 group, and the GGR-1 group (here renamed to LGC-57 group) (Figure 1B) and contains many channels whose activating ligand and function are unknown. To investigate the properties of these channels, we first expressed each channel gene in *Xenopus* oocytes and tested for current activation during the perfusion of a panel of neurotransmitters. Despite their homology to vertebrate GABA_A_ and glycine receptors (the source of the name GGR-1), we observed no activation of any members of the LGC-57 group (or any diverse group channels) by either GABA or glycine. Instead, we found three closely related channels of the LGC-57 group: LGC-57 (formerly GGR-1), LGC-58 (formerly GGR-2) and LGC-40 to be specifically gated by choline and acetylcholine (Figure 1A). All three channels showed a preference for choline, with EC_50_ values 2.5 to 3-fold lower for choline than acetylcholine (Figure 1C). These findings parallel a previous report that LGC-40 forms a choline and acetylcholine-gated channel, although in contrast to that report we did not observe serotonin responses (Ringstad et al., 2009). We did not observe currents in response to any of the tested compounds for the remaining members of the diverse group (Figure S1A). The lack of agonist induced currents may be because the channel was poorly expressed, the correct ligand was not tested or because they function only as components of heteromeric complexes. The remaining member of the LGC-57 group, LGC-39 showed unusual activation properties which will be discussed below.

**Figure 1.**
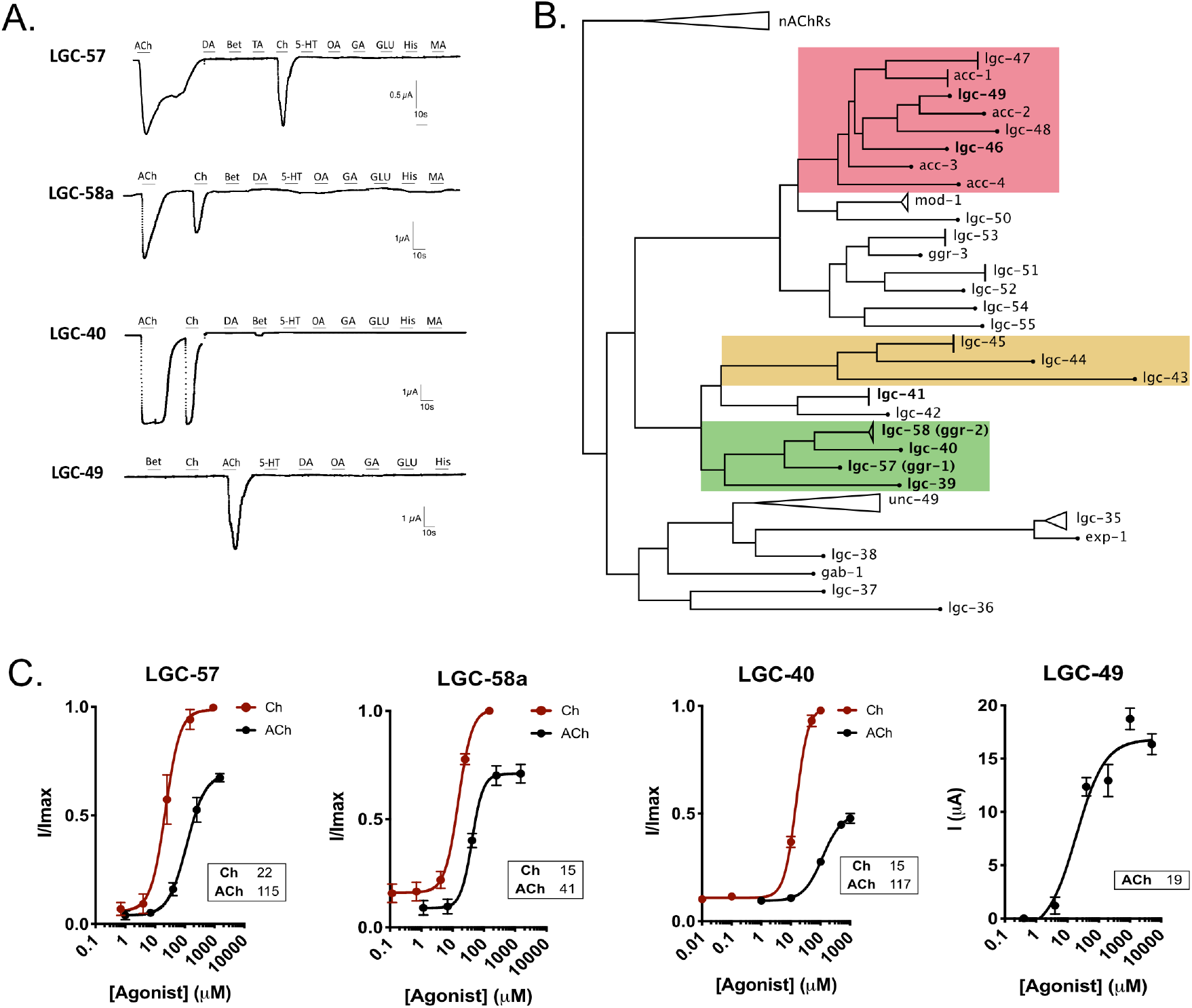
Deorphanisation of Cholinergic Ligand-Gated Ion Channels. (**A**) Continuous current traces of *Xenopus* oocytes expressing LGC-57, LGC-58a, LGC-40, and LGC-49, oocytes were perfused for 10s with a panel of ligands each at 1mM: ACh (acetylcholine), Ch (choline), Bet (betaine), TA (tyramine), DA (dopamine), 5-HT (serotonin), OA (octopamine), GA (GABA), GLU (glutamate), His (histamine), MA (melatonin). (**B**) Phylogenetic tree of a subgroup of Cys-loop ligand gated ion channels in *C. elegans*, groups of interest are highlighted by colour: ‘ACC’ group of acetylcholine gated channels (red), LGC-45 group (yellow), LGC-41 group (blue), LGC-57 group (green). (**C**) Dose response curves for LGC-57, LGC-58a, LGC-40 and LGC-49 in response to their major ligand(s). Current is normalised by I/Imax for each oocyte, for LGC-49 current is raw un-normalised current. Error bars represent SEM of 5-14 oocytes. Curves are fitted with a four-parameter variable slope, inserts show EC_50_ in µM for each ligand.

In addition to the diverse group channels, many other *C. elegans* LGICs lack identified ligands. For example, while several members of the ACC group (<u>A</u>cetylcholine-gated <u>C</u>hloride <u>C</u>hannels) of LGICs have been shown to form acetylcholine-gated chloride channels, four members of this subfamily *(acc-4, lgc-47, lgc-48,* and *lgc-49)* had not previously been characterised (Figure 1B, in red). Upon expression in *Xenopus* oocytes, we found that one of these channels, LGC-49, formed a homomeric acetylcholine-gated channel with an EC_50_ of 19µM (Figure 1C), similar to the EC_50_ values published for other members of this group (Putrenko et al., 2005; Takayanagi-Kiya et al., 2016). Unlike the members of the LGC-57 group, which showed activation by both acetylcholine and choline, LGC-49 showed no significant activation by choline. None of the ligands tested here induced currents for ACC-4, LGC-47 or LGC-48, either alone, or in combination with closely related and previously characterised LGICs, such as ACC-1 (Figure S1A&C) (Putrenko et al., 2005). Given the vast number of possible heteromeric combinations within the ACC group it may be that these orphan channels are part of more complex channel compositions not tested here.

We next investigated the ion selectivity of the newly-deorphanised channels by carrying out ion substitution experiments in oocytes expressing LGC-57, LGC-58, LGC-40, or LGC-49. For all these channels we observed significant reversal potential shifts following substitution of standard high NaCl buffer for low Cl (Na Gluconate), but not following substitution with Na^+^-free (NMDG) solution, indicating selectivity for anions over cations for all the tested channels (Figure 2A-B). We also tested the previously deorphanised channel LGC-46 (Liu et al., 2017), which to date lacked ion selectivity data. This channel likewise showed reversal shifts characteristic of an anion selective channel (Figure 2A-B). Interestingly, all members of the LGC-57 group possess a PAR motif (Proline-Alanine-Arginine), located in the M2-3 intracellular loop (Figure S1B), which has been shown to impart anion selectivity to LGICs (Wotring et al., 2003). Although several uncharacterised members of the ACC group have sequences that diverge from the PAR motif, both LGC-49 and LGC-46 contain the PAR motif sequence (Figure S1B). Thus, the PAR motif appears to correlate with anion selectivity in both the LGC-57 and ACC groups of nematode acetylcholine-gated LGICs.

**Figure 2.**
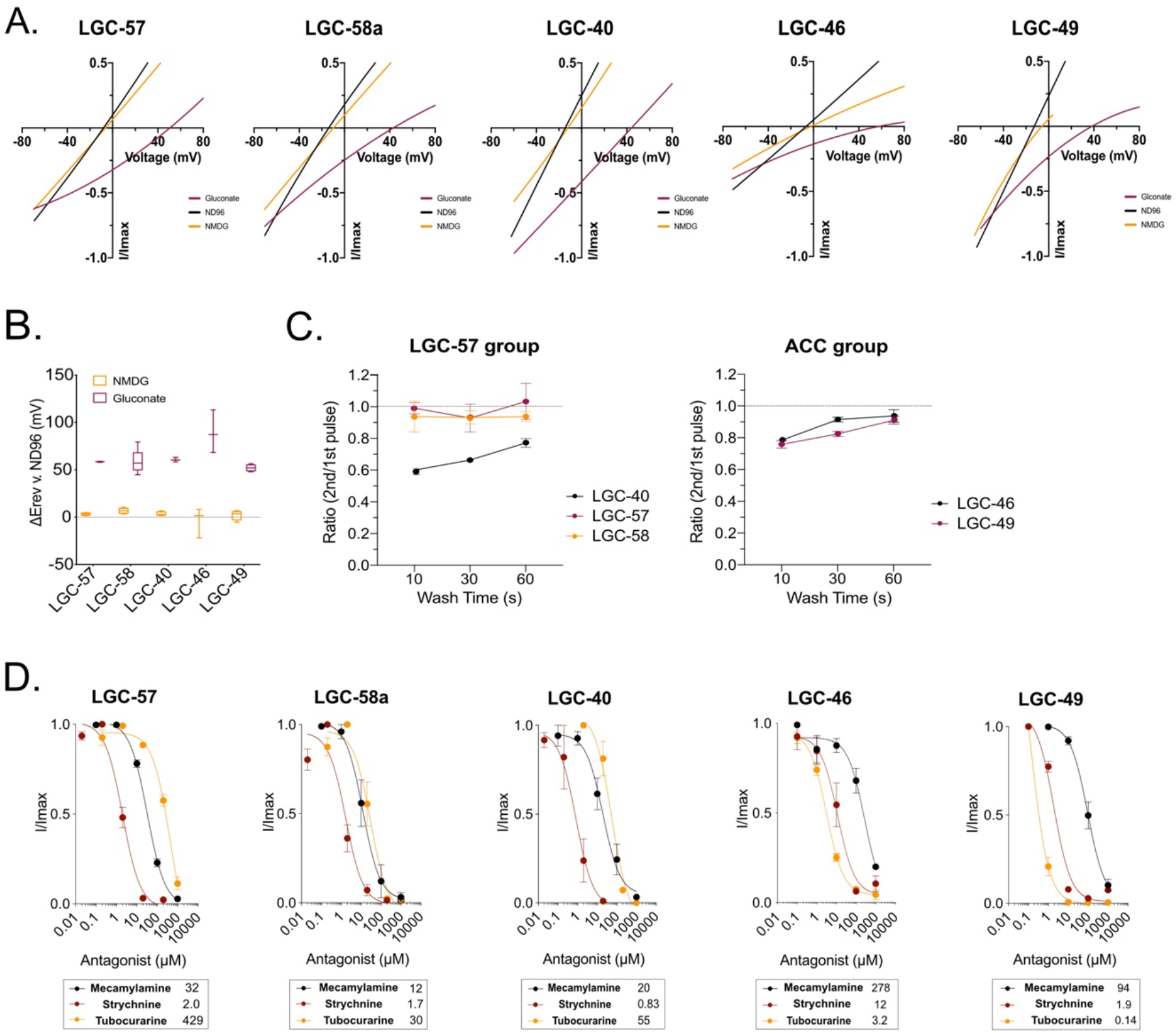
Ion Selectivity and Antagonistic Characterisation of Cholinergic LGICs. **(A)** Representative current voltage plots of newly deorphanised channels in ND96, Na gluconate or NMDG solutions. Current was normalised by leak current subtraction (in absence of activating ligand) and the peak current for each oocyte. (**B**) Box plot of ΔE_rev_ of NMDG and Na Gluconate vs. ND96 in oocytes expressing LGC-57, LGC-58a, LGC-40, LGC-39, LGC-46, and LGC-49, E_rev_ was calculated in the presence of the major agonist of each channel and leak subtracted. N=6-11 oocytes. (**C**) Current ratio of oocytes expressing LGC-57, LGC-58, LGC-46, LGC-49 or LGC-39 undergoing repeated agonist application with 10s, 30s and 60s wash intervals. Error bars represent SEM. N=5-9 oocytes per condition. (**D**) Antagonist application in the presence of ligands using mecamylamine, strychnine and tubocurarine. Current was normalised by I/Imax for each oocyte, curves are fitted with a three-parameter variable slope, error bars represent SEM of 2-7 oocytes, inserts show IC_50_ in µM for each antagonist.

### Cholinergic Channels Display Diverse Antagonist Binding Properties

To understand if there are further functional differences between the channels deorphanised in this study, we exposed each channel to three cholinergic antagonists, mecamylamine, strychnine and d-tubocurarine. Strychnine and d-tubocurarine have been shown to compete with the full agonist for the ligand binding domain, although their binding mechanisms vary between LGICs of different classes (Brams et al., 2011); in contrast, mecamylamine has been shown to interact with the transmembrane regions of mammalian nAChRs (Bondarenko et al., 2014). Indeed, we saw that the antagonistic profile differed significantly between the newly-deorphanised channels. For example, within the LGC-57 group the two smallest antagonists, mecamylamine and strychnine, had similar IC_50_ values for LGC-57, LGC-58a and LGC-40 (Figure 2D). However, tubocurarine, the largest molecule of the antagonists, displayed an 11-fold shift in IC_50_ for LGC-57 compared to LGC-58 and LGC-40 (Figure 2D). Thus, the binding capabilities of tubocurarine on LGC-57 differs substantially from that of its closest family members LGC-58 and LGC-40. Likewise, in the ACC group, LGC-46 and LGC-49 could both be blocked by mecamylamine, strychnine and tubocurarine (Figure 2D). These dissimilarities again highlight the discrete differences between channels from the same subfamily, which may have similar ligand-binding profiles for endogenous ligands. Interestingly, tubocurarine was the most potent blocker for the ACC group channels LGC-46 and LGC-49, whereas this antagonist was the least effective of the channels tested in the LGC-57 group.

We also tested the channels’ responses to repeated stimulation by their primary ligand. We found LGC-40 to be sensitive to repeated stimulation, displaying a significant difference in ratio between the first and second pulse after both 10s of and 60s of washing intervals (Figure 2C). In contrast, all other channels were capable of fast activation intervals as they did not display any decrease in peak amplitude after repeated stimulation (Figure 2C). Thus, LGC-40 appears to desensitise more than the other channels following activation by its ligand.

### LGC-39 is a Novel Polymodal Channel Activated by Cholinergic and Aminergic Ligands

One channel from the LGC-57 group, LGC-39, showed distinct ligand binding properties from the rest of the group. Unlike the other LGC-57 subfamily members, LGC-39 showed relatively little activation by choline (Figure 3A). Moreover, while acetylcholine activated LGC-39 strongly (with an EC_50_ of 1290µM), the most potent ligands for LGC-39 were the monoamines octopamine and tyramine (with EC_50_ values of 921µM and 686µM respectively; Figure 3A-B). In addition, LGC-39 also displayed small currents in response to dopamine (Figure 3A). Both activation by aminergic or cholinergic ligands resulted in a sigmoidal shaped dose response curve, which would indicate that all ligands bind in a similar positive cooperative manner in the same pocket (Cattoni et al., 2015). Like other members of the LGC-57 family, LGC-39 contains the PAR sequence (Figure S1B), and LGC-39-expressing oocytes showed reversal potential shifts in response to chloride but not sodium substitution (Figure 3C). Thus, *lgc-39* appears to encode a homomeric anionic and polymodal channel, capable of being activated by both aminergic and cholinergic neurotransmitters (Figure 3A-C).

**Figure 3.**
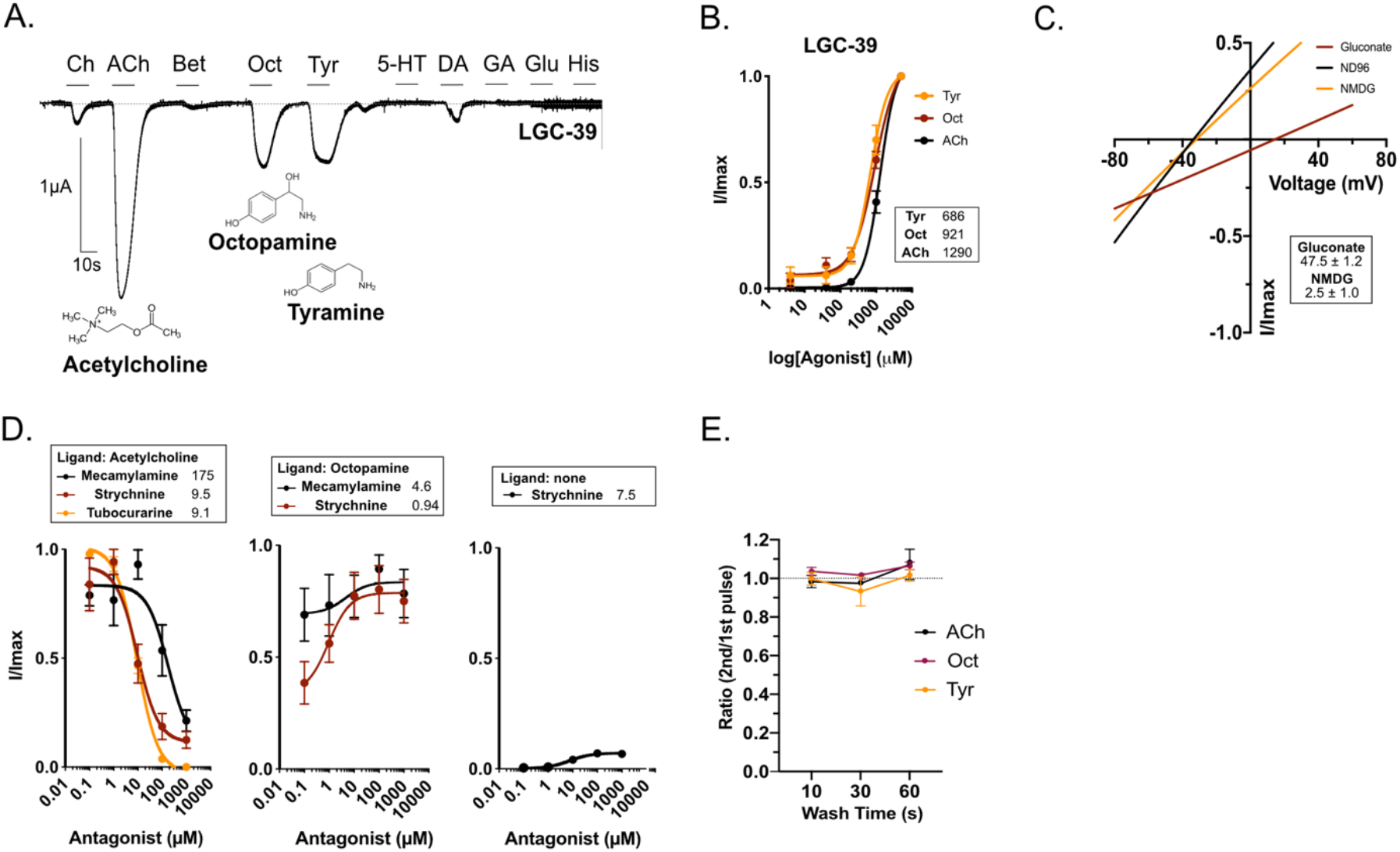
LGC-39 Forms a Polymodal Ligand-Gated Ion Channel. (**A**) Continuous current trace of a *Xenopus* oocyte clamped at -60mV expressing LGC-39, perfused during 10s pulses with a panel of ligands each at 1mM: ACh (acetylcholine), Ch (choline), Bet (betaine), Tyr (tyramine), DA (dopamine), 5-HT (serotonin), Oct (octopamine), GA (GABA), Glu (glutamate), His (histamine). (**B**) Dose response curve for LGC-39 in response to ACh, Oct and Tyr. Current is normalised by I/Imax for each oocyte. Error bars represent SEM of 8-12. Curves are fitted using a four-parameter variable slope, inserts show EC_50_ in µM for each ligand. (**C**) Current-voltage relationship during recordings in NMDG, Na Gluconate or ND96 in oocytes expressing LGC-39. Insert shows ΔE_rev_ vs. ND96 in mV +/- SEM of 5 oocytes. (**D**) Antagonist dose response curves for LGC-39 expressing oocytes activated by either ACh, Oct or no ligand at a constant dose and varying the antagonist doses (mecamylamine, strychnine and tubocurarine). Error bars represent SEM of 3-7 oocytes. Curves are fit with a three-parameter variable slope, inserts show IC_50_ in µM for each ligand. (**E**) Three different agonists do not differ in how they influence the ability for LGC-39 to be stimulated with short time intervals after repeated stimulation.

We tested the effects of antagonists on LGC-39 currents evoked by different activating ligands. In the presence of acetylcholine, LGC-39 could be blocked by the antagonists mecamylamine, strychnine and tubocurarine (Figure 3D). In contrast, the octopamine response could not be blocked by mecamylamine or strychnine. Surprisingly, strychnine, without the presence of an activating ligand, acted as a partial agonist, since it induced a small current with an EC_50_ of 7.5µM (Figure 3D). To further separate the functionality of the ligands, we also investigated if repeated activation by the different ligands influenced the ability for reactivation of LGC-39 differently (Figure 3E). No difference was seen for any wash interval between the ligands, which could suggest that all ligands occupy the binding site in a similar time frame or that the recovery time for the receptor is independent of the activating ligand.

### Cholinergic Channels show Broad and Varied Expression in The *C. elegans* Nervous System

To gain insight into the roles of cholinergic LGICs in the nervous system, we generated reporter lines to characterise their neural expression patterns. We used a similar set of fluorescent reporter lines to characterise the expression pattern of the newly deorphanised LGICs, by using transcriptional reporter transgenes in which the upstream promoter of the *lgc* gene drives the expression of a fluorescent protein. We then identified transgene-expression based upon location, morphology and known marker lines. Using such a transcriptional reporter, we observed primarily neuronal expression of the genes in the LGC-57 group, with little overlap observed in the neurons that were expressing reporters for *lgc-40*, *lgc-57* and *lgc-58* (Figure 4A-C). *lgc-40* was expressed in many pharyngeal neurons (M2, M3, MC, MI, I2), *lgc-57* in the A-class and B-class motorneurons of the ventral cord, and *lgc-58* in the egg-laying motorneurons (VCs; *lgc-57* was also observed in a subset of VCs). This suggests that these channels are likely to exist primarily as homomers *in vivo* and function in distinct target neurons. Further we observed the reporter for *lgc-39* in a range of interneurons and motor neurons, including the AVA premotor interneurons (Figure 4E). In addition to receiving extensive cholinergic input, the AVA neurons are the major synaptic target for the only octopaminergic neurons, the RICs, suggesting that LGC-39 may be exposed to both octopamine and acetylcholine *in vivo* (Figure 4F) and may be involved in both cholinergic and octopaminergic synaptic transmission. Finally, we found that the ACC group channel, *lgc-49,* was expressed in sensory neurons, including posterior sensory neurons such as ALN and PLN (Figure 4D).

**Figure 4.**
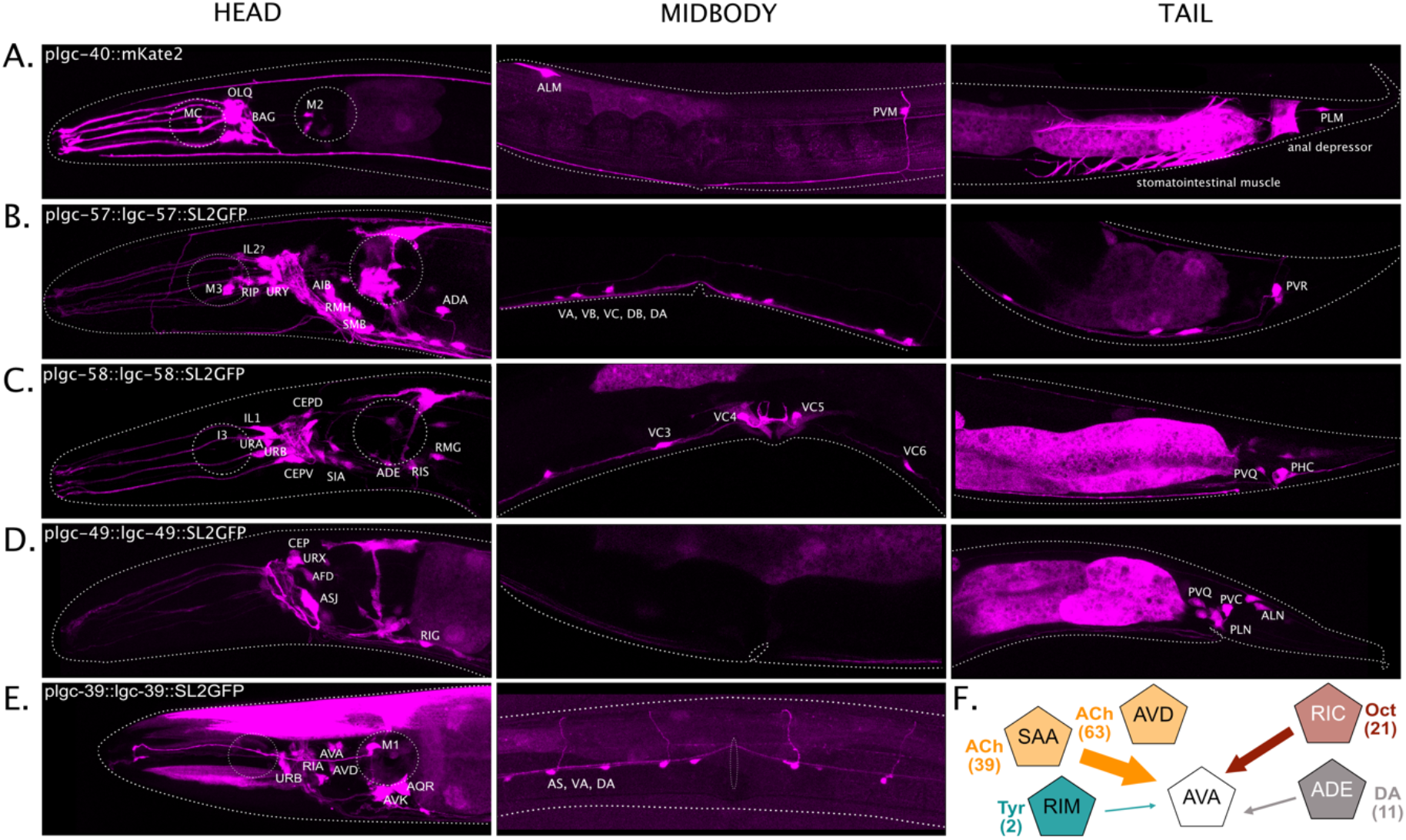
Newly Deorphanised LGICs are Expressed Broadly in the Nervous System. **(A-E)** Fluorescent reporters of intercistonically spliced mKate2 or GFP driven under the promoter and/or genomic sequence of reveals broad neuronal expression of cholinergic channels with little overlap. (**F**) Schematic depicting a subset of the synaptic connections received by the *lgc-39* expressing neuron class, AVA, numbers in brackets show the total number of synapses for each connection.

We also used reporters to analyse the expression pattern of several still-orphan LGICs, from the diverse and ACC groups, including *lgc-42, lgc-47, lgc-48, lgc-43* and *lgc-45*, as well as the previously deorphanised acetylcholine-gated channel *lgc-46* (Takayanagi-Kiya et al., 2016) (Figure S2A-D). These reporters also showed diverse and distinct patterns of expression, primarily in neurons. For example, *lgc-46* was broadly expressed in several neurons, mostly in the head (Figure S2D). Most of the orphan channels were also expressed specifically in neurons; for *lgc-47* this expression was unusually broad, encompassing sensory, motor, and interneurons (Figure S2A), while *lgc-48* was expressed only in a single pair of neurons, the ADL chemosensory neurons (Figure S2B). Interestingly, the two orphan channels, *lgc-43,* and *lgc-45,* which lack a PAR sequence and may thus encode cationic channels (Figure S1B), did not appear to be expressed in neuronal tissue, but instead in the hypodermis (Figure S2B). Together, these data suggest that these channels play various roles in, and outside, the nervous system.

### Excitatory and Inhibitory ACh Receptors are Co-expressed in Many Neurons

Our fluorescent reporter expression analysis indicated that many of the newly deorphanised inhibitory ACh-gated receptors in this study are expressed in neurons previously shown to also express excitatory ACh-gated channels (Barbagallo et al., 2010; Raizen et al., 1995). These results imply that acetylcholine, as an inhibitory neurotransmitter, may have a larger role than previously appreciated, and that acetylcholine contributes to both inhibitory and excitatory events in many neurons. To determine the extent to which excitatory and inhibitory receptors, for the same neurotransmitter, are expressed in individual neural classes, we made use of the single cell RNAseq dataset from *C. elegans* neurons (Taylor et al., 2021). We first generated a complete list of ionotropic receptors for each of the three classical neurotransmitters ACh, GABA and glutamate (Table S1). Since receptors with unknown ligand-identity would have the potential to bias predictions, we predicted the ligand and ion selectivity of orphan channels based upon homology with closely related characterised channels, and the presence, or absence, of a PAR motif in the M2-3 intracellular loop (see Methods).

From this analysis, we found a remarkable frequency of neural classes that co-express both inhibitory and excitatory receptors for the same neurotransmitter. This was particularly notable for acetylcholine, for which over 60% of the neural classes expressed both excitatory and inhibitory ACh receptors. In contrast, GABA receptors were more biased toward inhibition, with over 40% of neural classes expressing only inhibitory GABA receptors (Figure 5A, S3). To make generalised predictions of synaptic polarity, we summed expression of inhibitory and excitatory receptors, for each neurotransmitter, in each neural class and assigned synapse polarity based on the most prevalent receptors in each neural class, assuming that all receptors in a cell are present equally at all synapses (Figure 5C, S4). This analysis suggested that the majority of ACh and glutamate synapses are excitatory and most GABA synapses are inhibitory, though this varied significantly for individual connections (Figure 5B).

**Figure 5.**
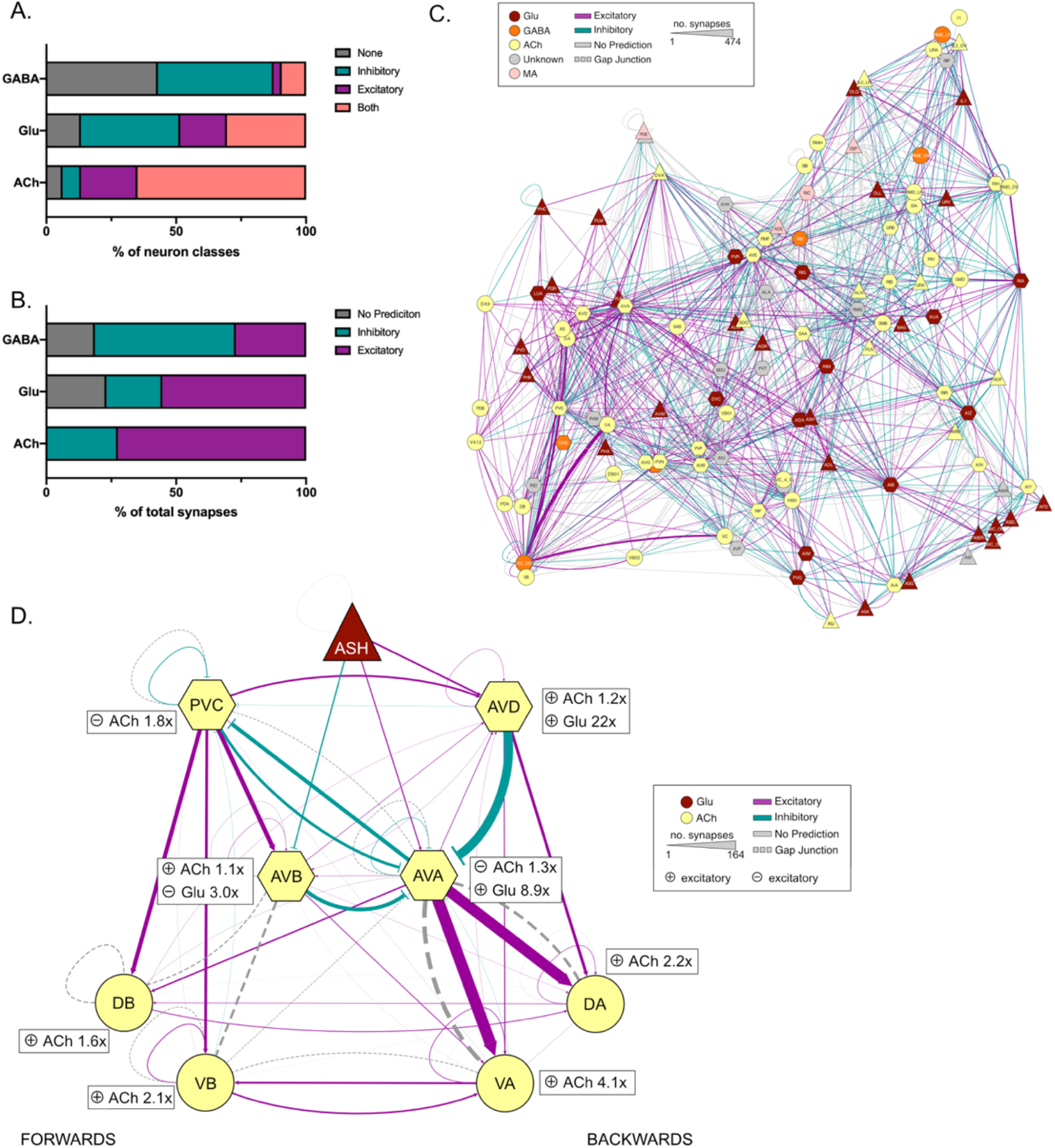
Predicting Synapse Polarity Based on LGIC Expression. (**A**) Bar chart depicting the percentage of total neuron classes expression inhibitory, excitatory, both or no receptors for GABA, glutamate (Glu) and acetylcholine (ACh). (**B**) Bar chart depicting the percentage of total synapses for a given neurotransmitter that are predicted to be inhibitory, excitatory or have no prediction. (**C**) Network diagram depicting the polarity of synaptic connections between neural classes, connection colour show polarity: teal (inhibitory), pink (excitatory), grey (no prediction). Gap junctions are represented in dashed lines. Line weight represents number of synapses and nodes are coloured by the major neurotransmitter a class release. Diagram made with cytoscape using the EntOpt clustering package. (**D**) Network diagram depicting the predicted polarity of the locomotion circuit, connection colour shows polarity: teal (inhibitory), pink (excitatory), grey (no prediction). Inserts next to each neuron node show the fold magnitude of expression of the major receptor type for each neurotransmitter in each neural class, e.g., AVA neurons express 1.3x as many inhibitory ACh receptors than excitatory and 8.9x as many excitatory glutamate receptors than inhibitory. Gap junctions are represented by dashed lines. Line weight represents number of synapses and nodes are coloured by the major neurotransmitter a class release. Diagram made with cytoscape.

To examine the validity of our polarity predictions we investigated the sign prediction using previously characterised neuronal circuits. We picked the well-studied locomotion circuit (Chalfie et al., 1985) consisting of the interneurons AVD, AVE and AVA, which initiate reversals, and PVC and AVB that initiate forward movement. Most of our predicted connection polarities (Figure 5D) were consistent with circuit data from previous studies, such as the excitatory connection between AVA and the VA and DA motor neurons, which is involved in controlling reverse locomotion, as well as the excitatory connection from the sensory neuron ASH to the reverse command neuron AVA (Mellem et al., 2002; Piggott et al., 2011). We also observed connections which appeared counter intuitive such as an inhibitory acetylcholine connection from AVD to AVA, two interneuron pairs thought to be co-ordinately active during reverse locomotion (Faumont et al., 2012) (Figure 5D). While some studies have proposed additional inhibitory connections within this circuit (Rakowski and Karbowski, 2017), AVA neurons express several ACh-gated channels and has a relatively low ratio of inhibitory to excitatory receptor expression (1:3), upon which this prediction was made. This suggests that some connections may indeed be both inhibitory and excitatory, especially where a neuron expresses a large range of different channels and receives input from many different neural classes. Connections such as these require further *in vivo* investigations to address these predictions.

### Determining Synaptic Localisation of LGICs

We reasoned that the single cell RNAseq dataset (Taylor et al., 2021) might also be useful for predicting the intracellular localisation of cholinergic LGICs, as presynaptic receptors would be predicted to be expressed in cholinergic neurons, while postsynaptic receptors should be expressed in neurons receiving cholinergic input. To assess the correlation between the number of cholinergic synapses a neuron makes (‘outgoing ACh synapses’) or receives (‘incoming ACh synapses’), with the expression level of cholinergic LGICs, we produced two heatmaps showing the expression of ACh-gated receptors, with neural classes ranked by the total number of incoming or outgoing ACh synapses (Figure 6A-B). This analysis highlights that the expression of some ACC group channels, in particular, *lgc-46,* correlate with both the number of incoming and outgoing ACh synapses (Figure 6C-D & Figure S5). This correlation suggests that these receptors may be acting either pre- or post-synaptically. In contrast, members of the choline and ACh-gated LGC-57 group, *lgc-57*, *lgc-58* and *lgc-40*, showed little correlation with either incoming or outgoing synapses (Figure 6C-D). Surprisingly for this subgroup, several cells with high ACh connectivity showed low receptor expression level (Figure 6C-D). This may be suggestive of an extrasynaptic role for these receptors, however further evidence is required to make these assumptions.

**Figure 6.**
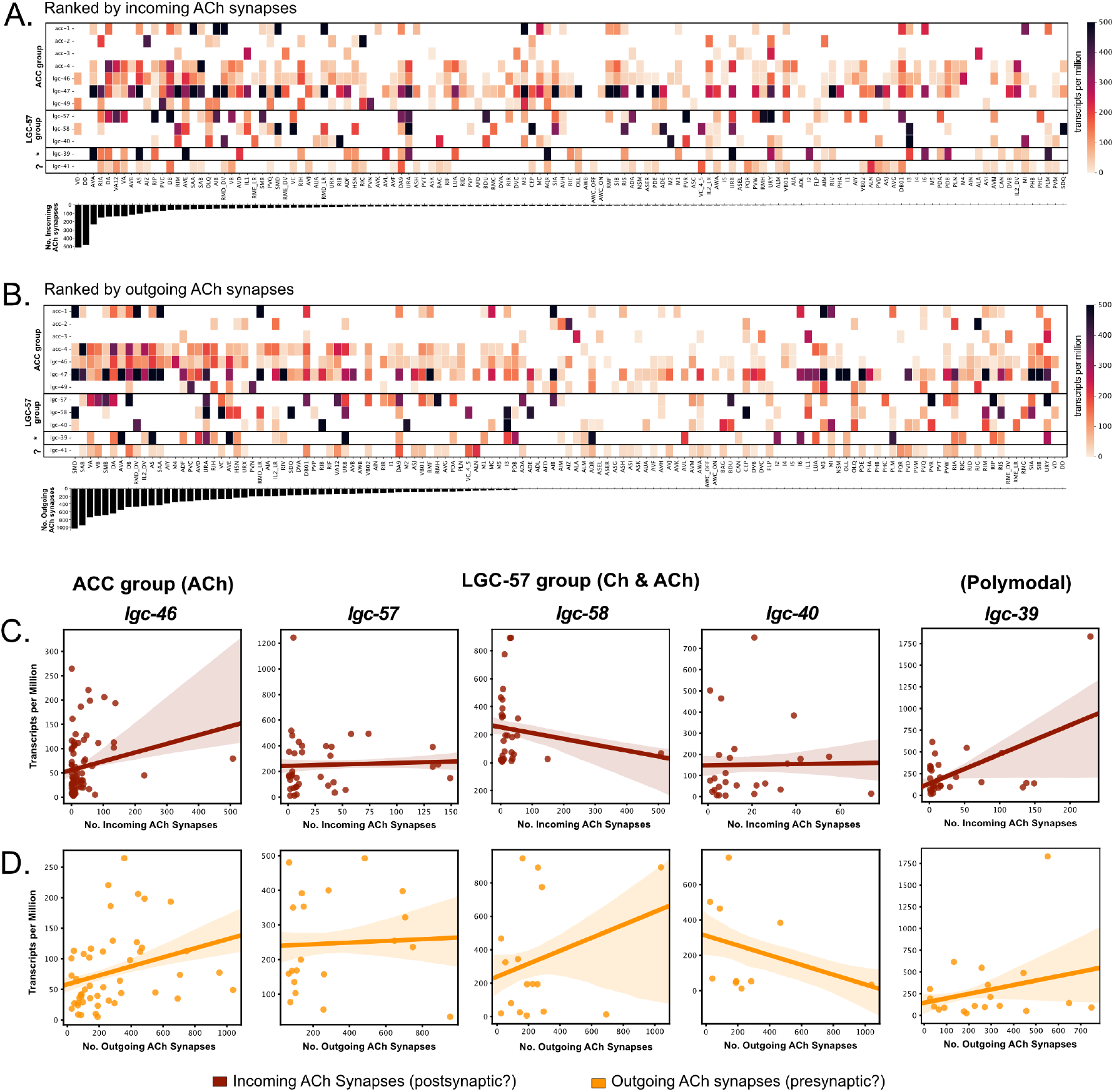
Correlation of Cholinergic Synapses with Expression Pattern of Cholinergic Ion Channels. (**A-B**) Heatmaps showing the expression level of newly deorphanised LGICs in each neural class. Neurons are sorted by the total number of cholinergic synapses they receive (top, ‘Incoming’) or make (bottom, ‘Outgoing’). (**C & D**) Scatter plots showing correlation between the total number of incoming (red) or outgoing (orange) cholinergic synapses for a given neuronal class and expression of *lgc-46, lgc-57, lgc-58, lgc-40,* and *lgc-39*.

To empirically assess the synaptic localisation of cholinergic LGICs, we generated endogenous GFP-tagged CRISPR lines for members of the LGC-57 subgroup, including *lgc-39*, *lgc-40, lgc-57,* and *lgc-58* and (Figure 7A-D). In all cases GFP was inserted in the intracellular M3/4 loop and the function of the resulting chimeric protein was verified in *Xenopus* oocytes (Figure S6A). We observed a clear difference in the localisation pattern for these channels. LGC-39::GFP was localised in distinct punctate structures both in the nerve ring and along the ventral cord, suggestive of synaptic localisation (Figure 7A), and consistent with the positive correlation between *lgc-39* expression and incoming and outgoing ACh synapses (Figure 6A-D). Members of the choline-gated LGC-57 group however showed diffuse protein expression. LGC-40::GFP appeared to have diffuse expression in the nerve ring, and touch receptor neurons, with cell bodies often being visible (Figure 7B). Notably, cell body LGC-40::GFP expression was detected in the posterior and anterior bulbs, in cells which anatomically correspond to MC and M2 neurons (Figure 7B). While LGC-57::GFP appeared to have overall low expression and little protein localisation could be seen above background (Figure 7C). LGC-58::GFP was clearly visible in the nerve ring and VC4/5, including some punctate structures (Figure 7D). Since these choline-sensitive members of the LGC-57 group showed little correlation with ACh synapses, their diffuse protein localisation may be indicative of an extrasynaptic role (Figure 6C & Figure 7D).

**Figure 7.**
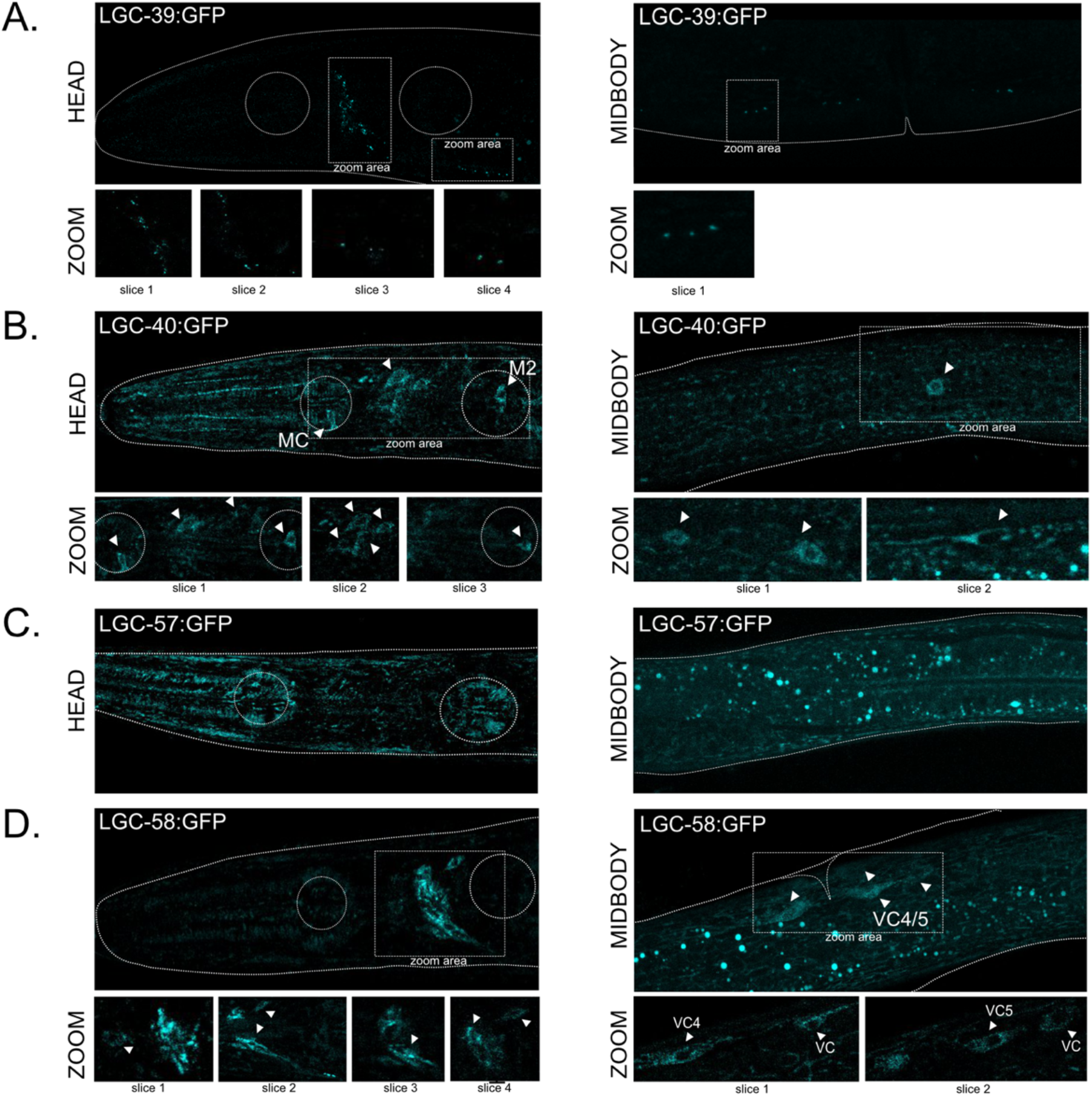
Protein expression pattern of cholinergic ion channels. (**A-D**) Localisation of endogenously GFP tagged LGC-39, LGC-40, LGC-57 and LGC-58. White arrows highlight areas of interest, represented in higher magnification below.

## Discussion

### A Novel Family of Cholinergic LGICs

This study highlights the diversity among cholinergic LGICs in *C. elegans*. Nematodes have previously been shown to express acetylcholine and choline-gated excitatory LGICs related to nicotinic receptors, as well as inhibitory acetylcholine-gated chloride in the ACC group. Here we describe a second inhibitory subfamily, that contains channels gated by both choline and acetylcholine: LGC-57, LGC-58 (previously named GGR-1 and GGR-2) and LGC-40. In contrast to the ACC group of acetylcholine-gated anion channels, these newly deorphanised channels are gated preferentially by choline, the metabolite of acetylcholine which is abundant at cholinergic synapses.

These results add to the already extensive catalogue of acetylcholine-gated channels in *C. elegans* (Putrenko et al., 2005; Takayanagi-Kiya et al., 2016), and to the growing number of choline-gated channels described in *C. elegans,* which previously consisted of the excitatory DEG-3/DES-2 channel found within the nAChR superfamily (Yassin et al., 2001). Together with our new data this highlights the expansion and importance of cholinergic transmission in nematodes. These newly deorphanised channels display subtle variations in their ability to bind ligands and antagonists, which translates into physiologically relevant differences that may increase the fine tuning in the control of neuronal transmission and contribute to complex neuronal signalling within a relatively minimal neuronal network. Interestingly, the electrophysiologically similar channels LGC-57, LGC-58, and LGC-40 show largely distinct patterns of expression within the nervous system of *C. elegans*, suggesting they may form homomeric channels with distinct functions *in vivo*. When tagged with a fluorescent protein, these three channels also showed a diffuse localisation pattern within the neuron, suggesting that in contrast to the ACC group channels, such as LGC-46 (Takayanagi-Kiya et al., 2016), these channels may not be synaptically localised. This suggests a possible distinct extrasynaptic role for choline by acting via these channels, in the modulation of the nervous system.

The observation that choline shows higher efficacy for these channels, a molecule generated at cholinergic synapses through catabolism of acetylcholine by cholinesterases, raises the possibility that choline is their true *in vivo* ligand and that choline itself may function as a neuromodulator. The idea that choline could activate cholinergic receptors differently from acetylcholine has been discussed for other cholinergic receptors that can be dose-dependently blocked or activated by choline (Purohit and Grosman, 2006). Here we have identified cholinergic receptors in which choline act as a full agonist, showing preference in binding and activation degree for choline over acetylcholine. Previous reports suggest aromatic residues in the extracellular domain of mammalian neuromuscular AChRs play a vital part in stabilising the binding of acetylcholine over the binding to choline (Bruhova and Auerbach, 2017). Interestingly, at least some of these aromatic residues are lacking in the choline-gated receptors identified here. Thus, it is not unreasonable to hypothesise that choline could be an authentic endogenous ligand for these channels in *vivo*.

### A Polymodal LGIC Activated by Both Aminergic and Cholinergic Ligands

In this study we also identified a novel polymodal channel, LGC-39, which was gated not only by acetylcholine, but also by the aminergic ligands tyramine and octopamine. We observed dose-dependent activation of LGC-39 channels by these structurally distinct ligand classes of endogenous ligands at similar, physiologically-relevant concentrations. The expression pattern of *lgc-39* suggests that the channel might be exposed to all these ligands *in vivo*; for example, *lgc-39* is highly expressed in the AVA premotor neurons, which receive a large amount of input from acetylcholine producing neurons (White, 1986), as well as from the RIC neurons, the only octopamine producing cells in the *C. elegans* nervous system (Alkema et al., 2005). The AVA neurons also receive some input from tyraminergic and dopaminergic neurons, transmitters which we also found can activate LGC-39. Interestingly, in contrast to the choline-gated channels LGC-40, LGC-57, and LGC-58, we observe clear punctate localisation of LGC-39 in both the nerve ring and along the ventral cord, with no fluorescence visible in the cell bodies. This may suggest a role for LGC-39 as postsynaptic receptor for both cholinergic and aminergic neurotransmission.

The concept of a truly polymodal receptor, that can be activated by structurally diverse compounds, has not before been investigated in great detail, though previous observations have described roles for receptors that can use dual ligands for allosteric modulation (Cummings and Popescu, 2015). For example, dopamine exhibits a pseudo competitive ability to antagonise GABA_A_ currents, although this effect cannot be blocked by competitive GABA_A_ antagonists, which bind the main binding pocket (Hoerbelt et al., 2015). Further, D-serine has been shown to function as an allosteric modulator of NMDA receptor activity (Wolosker and Balu, 2020). In contrast to these examples, based on their capability to achieve dose-dependent activation by both amines and acetylcholine, both groups of neurotransmitters appear to be true ligands of LGC-39, most likely interacting with the ligand-binding domain. Understanding the mechanisms by which these multiple neurotransmitters can activate LGC-39 and potentially affect different behavioural outputs will be of interest in future studies.

### Functional Insights into The *C. elegans* Connectome

With increasing molecular and physiological characterisation of neurotransmitter receptors in *C. elegans*, it is becoming feasible to more accurately predict the functionality of synapses in the *C. elegans* connectome. In this study we used the expression pattern of newly and previously deorphanised LGICs for the three classical neurotransmitters, ACh, glutamate and GABA, to predict the polarity of synapses in the *C. elegans* connectome. By assigning synapse polarity based on relative expression levels of anionic and cationic receptors, we have provisionally predicted the sign of chemical synapses involving classical neurotransmitters. Although similar attempts to assign polarity to *C. elegans* synapses have been made in the past (Fenyves et al., 2020), these predictions were based upon incomplete or incorrect ligand assignment for many LGICs. These revised predictions correlate well with experimental data for many well-characterised circuits, such as the excitatory connections between the ASH nociceptors and the AVA interneurons, as well as between the AVAs and the VA and DA motorneurons (Mellem et al., 2002; Piggott et al., 2011). Our predicted inhibitory connection between AVB and AVA interneurons also correspond well with empirical data on the locomotor circuit (Kawano et al., 2011; Qi et al., 2012). In addition to providing sign predictions for synaptic connections, our model also provides a ratio of excitatory to inhibitory expression for each neuronal class and neurotransmitter. Not only do our predictions generate interesting functional hypotheses for future investigation, but this additional information also allows these predictions to be critically assessed. It also raises the question whether these connections, with low receptor ratios, represent truly complex connections, a question that could be addressed in future studies.

A surprising outcome of the expression analysis was the high frequency with which individual neurons expressed cationic and anionic receptors for the same neurotransmitter. This was especially prevalent for acetylcholine; our analysis indicated that 60% of neural classes express both inhibitory and excitatory receptors for acetylcholine, 30% for glutamate and 10% for GABA. One explanation for this apparent paradox is that excitatory and inhibitory receptors might be differentially localised in neurons, with some found extrasynaptically and others enriched in synapses. Various *C. elegans* LGICs are known to act in regions other than the post-synapse; for example, *lgc-35* has been shown to mediate GABA spill-over transmission (Jobson et al., 2015), while *lgc-46* appears be localised to the pre-synapse as an autoreceptor (Takayanagi-Kiya et al., 2016). The choline-sensitive channels from the LGC-57 group likewise appear to be extrasynaptic in their localisation (Figure 7). In addition, postsynaptic sites might themselves contain a mixture of excitatory and inhibitory receptors, which could differ in their ligand affinity, desensitisation kinetics and regulation. Indeed, LGIC localisation is not static; for example, glutamatergic AMPA receptors have been shown in many species to increase their synaptic localisation during learning (Malinow and Malenka, 2002), and recent evidence indicates that *C. elegans* LGICs also display regulated membrane trafficking upon learning (Morud et al., 2021). Thus, *C. elegans* may contain large numbers of complex cholinergic synapses with the potential to be excitatory or inhibitory depending on context or experience.

## Materials and Methods

### *C. elegans* culture

Unless otherwise specified, *C. elegans* worms were cultured on NGM agar plates with OP50 (Stiernagle, 2006). A full list of strains used in this study can be found in table S1.

### *Xenopus laevis* oocytes

Defolliculated *Xenopus laevis* oocytes were obtained from EcoCyte Bioscience (Dortmund, Germany) and in ND96 (in mM: 96 NaCl, 1 MgCl_2_, 5 HEPES, 1.8 CaCl_2_, 2 KCl) solution at 16° C for 3-7 days.

### Molecular biology

Unless otherwise specified, cDNA sequences of *C. elegans* genes were cloned from wildtype N2 worm cDNA (generated by reverse transcription PCR from total worm RNA using Q5 polymerase (New England Biosciences)). Where multiple isoforms are present isoform a was used. LGC-39 cDNA was generated by gene synthesis (ThermoFischer). For expression in *Xenopus* oocytes, ion channel cDNA sequences were cloned into the KSM vector downstream of a T3 promoter and between *Xenopus* β-globin 5’ and 3’UTR regions using the HiFi assembly protocol (New England Biosciences). *C. elegans* expression constructs were also generated using the HiFi assembly protocol (New England Biosciences) into the pDESTR4R3II backbone. *C. elegans* gDNA sequences were cloned from wildtype N2 gDNA and expression verified by the addition of GFP or mKate2 introduced on the same plasmid after an intercistronic splice site (SL2 site). Unless otherwise specified promoter sequences consist of approximately 2kb of gDNA upstream of the start codon.

### CRISPR/CAS9-mediated gene manipulation

Endogenous tagging of the M3/4 cytosolic loop of *C. elegans* LGIC proteins with GFP was carried out either using the SapTrap protocol (Dickinson et al., 2018; Schwartz and Jorgensen, 2016) for *lgc-39(lj121)*, or by SunyBiotech (Fuzhou, China) for *lgc-57(syb3536)*, *lgc-58(syb3562)*, and *lgc-40(syb3594)*.

### RNA synthesis and microinjection

CRNA was synthesised *in vitro* using the T3 mMessage mMachine transcription kit according to manufacturer’s protocol to include a 5’ cap (ThermoFischer Scientific). Prior to injection RNA was purified using the GeneJET RNA purification kit (Thermo Fischer Scientific). Size sorted and defolliculated *Xenopus* oocytes (Ecocyte) were placed individually into 96-well plates and injected with 50 nL of 500 ng/µL RNA using the Roboinject system (Multi Channel Systems GmbH). When two constructs were co-injected the total RNA concentration remained 500 ng/µL, with a 1:1 ratio of the components. Injected oocytes were incubated at 16°C in ND96 until the day of recording, typically between 3-6 days post injection.

### Two-Electrode Voltage Clamp (TEVC) recording and data analysis

Two-electrode voltage clamp recordings were carried out using either the Robocyte2 system or a manual set up with an OC-725D amplifier (Multi Channel Systems GmbH). Glass electrodes with a resistance ranging from 0.7-2 MΩ were pulled on a P1000 Micropipette Puller (Sutter). Electrodes contained AgCl wires and backfilled with a 1.5M KCl and 1 M acetic mixture. Unless otherwise stated, oocytes were clamped at -60mV. Continuous recordings at 500Hz were taken during application of a panel of agonists or antagonists. Data was recorded using the RoboCyte2 control software, or with WinWCP for manual recordings, and filtered at 10 Hz.

Dose response protocols used 10s agonist application pulses with 60s of wash in ND96 between each dose. Data was gathered over at least two occasions, using different batches of oocytes. Peak current for each dose was normalised to the oocyte maximum current using a custom-built python script (Morud et al., 2021). Normalised data was imported into Graphpad (Prism) and fitted to either a three or four parameter nonlinear Hill equation to obtain the highest degree of fit and calculate the EC_50_. Antagonist dose responses and ion selectivity recordings were carried out using the EC_50_ concentration of the primary agonist. Antagonist dose response protocols used 10s agonist + antagonist windows, with 60s of ND96 washes between doses. The agonist concentrations remained constant. Antagonist IC_50_ values were calculated using a second custom-built python script (Morud et al., 2021). Normalised data was imported into Graphpad (Prism) and fitted to either a three or four parameter nonlinear Hill equation to obtain the highest degree of fit and calculate the IC_50_.

Ion selectivity was detected using a voltage ramp protocol from -80mV to +60mV (20mV/s) in the presence of the primary agonist in three different solutions: ND96, NMDG (Na^+^ free) and Na Gluconate (low Cl^-^) solutions. Data was normalised to max current and ΔE_Rev_ was calculated using a custom-built python script (Morud et al., 2021). The resulting individual values or mean, SD and n for each construct was imported in GraphPad for further plotting and statistical analysis. Statistically significant differences in ΔE_Rev_ were calculated in GraphPad using a two-way ANOVA with Tukey’s correction for multiple comparisons. A representative normalised trace for each construct was also generated in Graphpad.

### Confocal and Cell ID

Worms were prepared and immobilised with 75 mM NaAzide in M9 and mounted onto 2% agarose in M9 pads. Image stacks were acquired with a 63x water immersion lens on a Leica SP8 or STED or using a 40x oil immersion objective on a Zeiss LSM780. Collapsed z-stack images were generated in Fiji/Image J. Neurons expressing fluorescent reporters were identified by cell shape, position and crossing with the multicolour reference worm NeuroPAL (Yemini et al., 2020).

### Synaptic polarity prediction

Inhibitory and excitatory chemical synapse prediction for ACh, Glu and GABA synapses were based upon expression levels of appropriate LGICs in postsynaptic cells. Chemical and electrical connectome data was obtained from Wormweb (http://wormweb.org/details.html), LGIC expression data was taken from the Cengen project using threshold level 4 (Taylor et al., 2021), ligand and ion selectivity for each channel was based upon this work. Previous work and predictions are presented in Table 1. Binary expression of LGICs for each neurotransmitter in each neural class were based upon expression and characterised in four groups: only excitatory, only inhibitory, both excitatory and inhibitory or none. These binary values were used to make the binary expression heatmap. Overall polarity of a synapse was calculated by summing the expression of all inhibitory and all excitatory LGICs for a given neurotransmitter in each cell class. The sum inhibitory was then taken from the sum of excitatory expression, resulting in an overall positive or negative signed expression in each neural class for each neurotransmitter. The ratio of these sums was also calculated to indicate the strength of polarity. It was assumed that each LGIC in each neural class is present equally at all synapses, therefore each incoming connection could be assigned a polarity based upon its receptor expression for that neurotransmitter. The resulting network with polarity was imported into cystoscope (Shannon et al., 2003) for plotting and further analysis. Analysis scripts can be found on GitHub at hiris25/Worm-Connectome-Polarity.

### Expression and cholinergic synapse analysis

The total number of cholinergic input or output synapses was calculated for each neural class by summing the number of presynapses for each cell that received a synapse from an ACh-producing neuron (incoming synapses), or the total number of post-synapses an ACh-producing neural class makes (outgoing synapses). Acetylcholine producing cells were described by (Pereira et al., 2015), the assumption was that all synapses made by an ACh-producing cell also release ACh, even when this cell co-transmits another neurotransmitter. Synapse number for each neuron was taken from (White, 1986). Expression data was obtained from (Taylor et al., 2021) using a threshold of 2. Neural classes were sorted by ACh in or out degree and the expression of each gene was mapped using a heatmap with an upper threshold of 500. For correlation plots, cells that did not express a receptor, were removed from the analysis. Correlation between expression level and ACh in, or out, degree was mapped using relplot in python’s seaborn package, confidence intervals were placed at 68%, corresponding to the standard error of the estimate.

## Data Availability

Python scripts can be found at on GitHub at hiris25/TEVC-analysis-scripts and hiris25/Worm-Connectome-Polarity. Aggregated data used for analysing TEVC data are available upon request from the Lead Contact. Further information and requests for *C. elegans* strains and plasmids is to be sent to and will be fulfilled by the Lead Contact William R Schafer

## Acknowledgements

The authors gratefully acknowledge Denise Walker and Lidia Ripoll Sanchez for help with cloning design, help with generating and maintaining strains, and other past and present members of the Schafer lab for helpful discussions. We would like to acknowledge the Centre for Cellular Imaging at the University of Gothenburg, Sweden, and the National Microscopy Infrastructure, NMI (NMI01125), for providing imaging facilities. This work was supported by grants from the Medical Research Council (MC-A023-5PB91), the Wellcome Trust (WT103784MA), National Institute of Health (W.R.S.), the Swedish Research council (VR2017-00236), Knut and Alice Wallenberg foundation (KAW2019,0293), Bollan stipend, Lundgrenska stiftelserna and Magnus Bergvalls Stiftelse (J.M.).

## Author Contributions

I.H., J.M. and W.R.S. designed the experiments. I.H., J.M., performed experiments and analysed data. I.H., J.M. and W.R.S. wrote the manuscript and all authors read and critically revised the manuscript to its final form.

## Supplementary Tables and Figure with Legends

**Supplementary Table 1:**
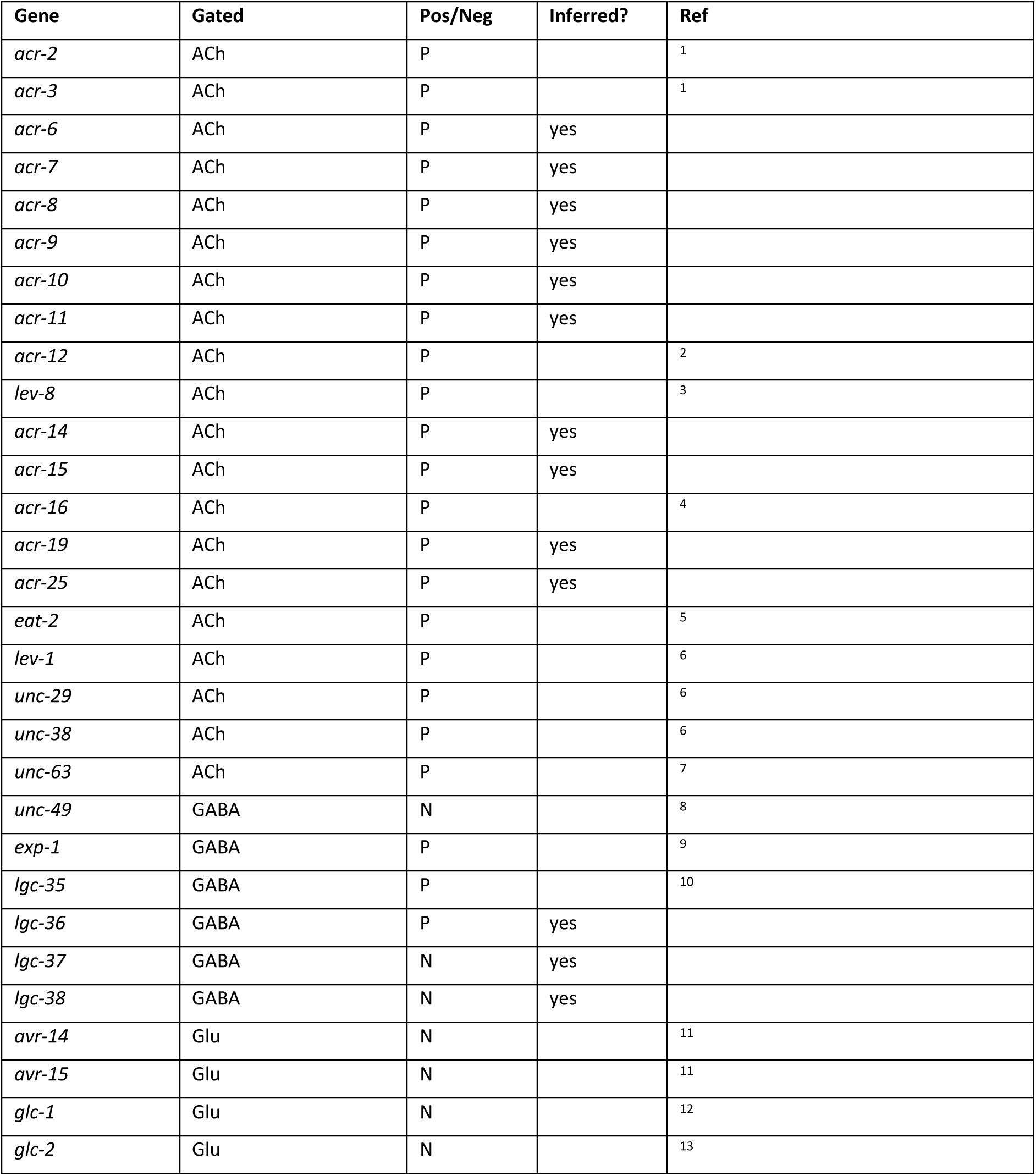
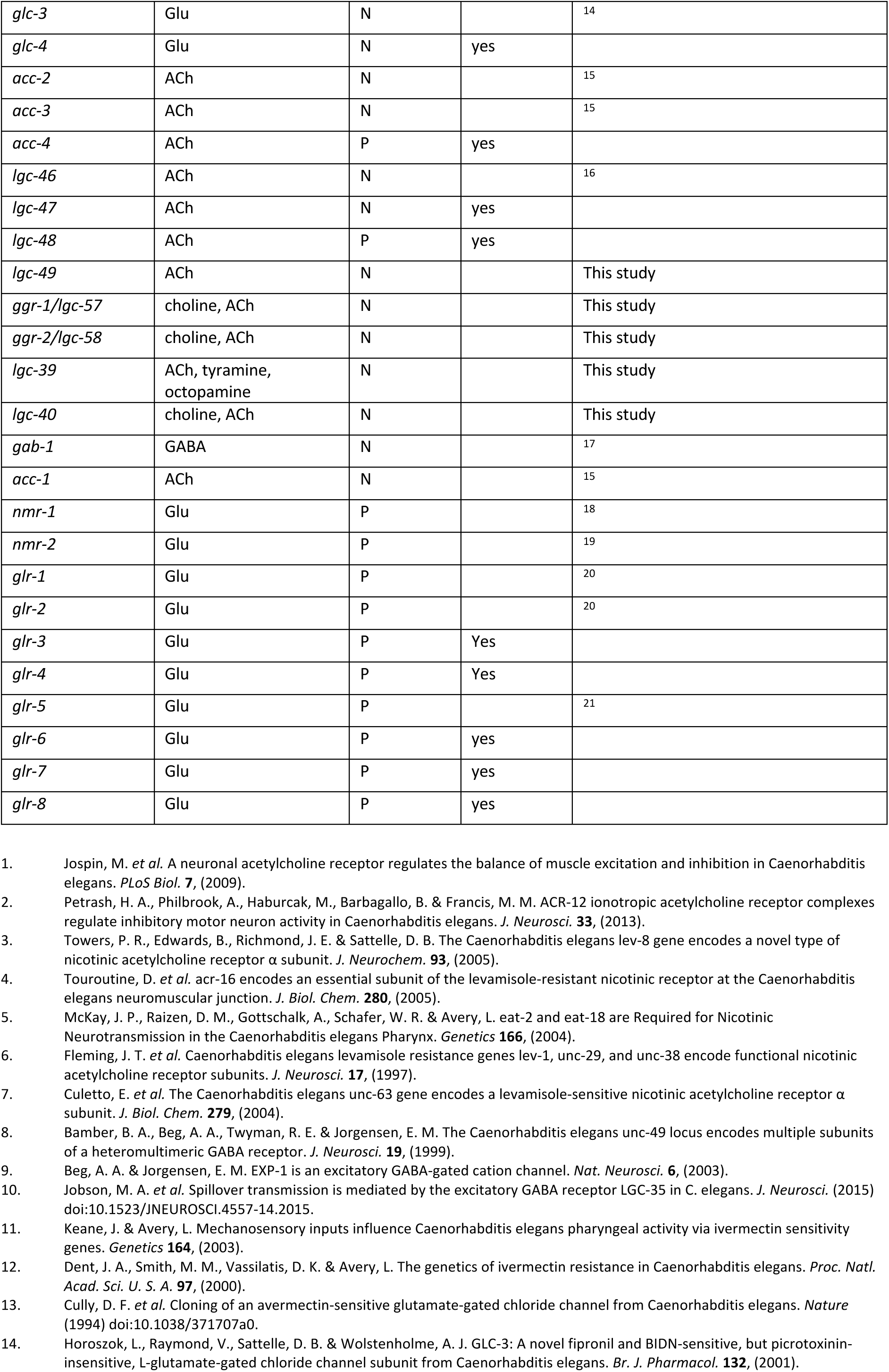
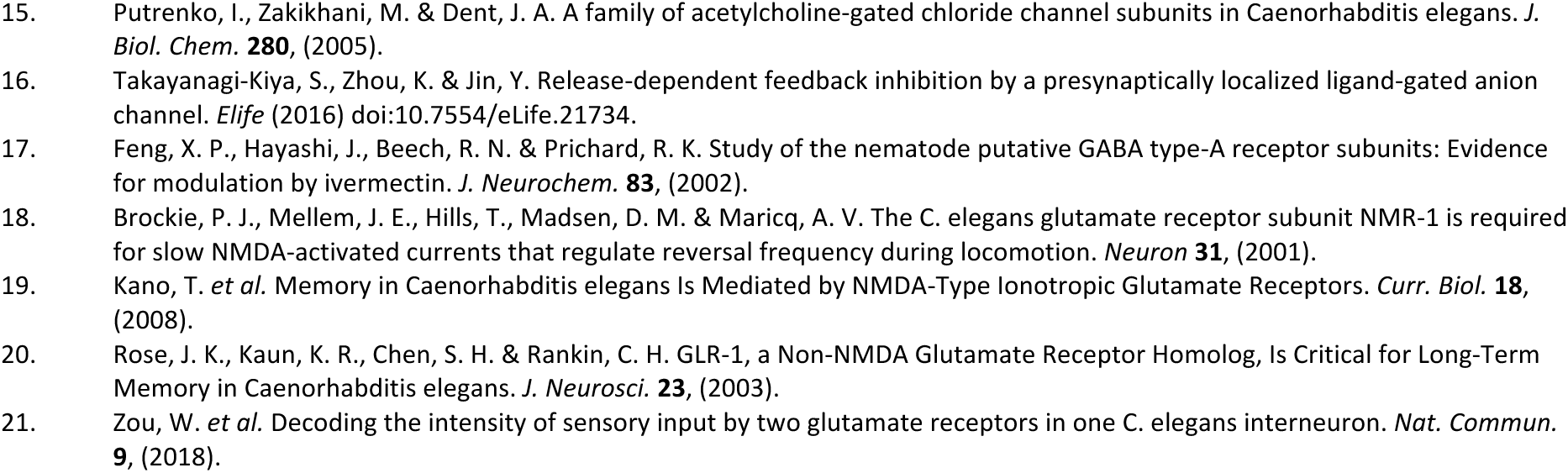
The list shows ligand identity and ion selectivity for LGICs in *C. elegans*. Each gene is assigned a polarity based upon its ion selectivity, this is shown in column ‘Pos/Neg’ as P: positive (cation selectivity) or N: negative (anion selectivity). For orphan receptors the ligand and ion selectivity has been predicted based on homology, this has been noted in the column ‘Inferred’ as ‘yes’.

**Supplementary Table 2:** List of *C. elegans* strains used in this study.

| Strain No. | Description |
| --- | --- |
| AQ4728 | him-5; lJEx1388[plgc-57::lgc-57::SL2GFP; punc-122::RFP] |
| PHX3594 | lgc-40(syb3594) |
| PHX3536 | lgc-57(syb3536) |
| PHX3562 | lgc-58(syb3562) |
| AQ4614 | lJEx1332[plgc-39(2kb)::lgc-39::SL2 GFP; punc-122::RFP] |
| AQ4427 | lJEx1254[plgc-40(2kb)::mKate2::gpd-23'UTR; unc-122::gfp] |
| AQ4423 | lJEx1252[plgc-57(2kb)::mKate2::gpd-23'UTR; unc-122::gfp] |
| AQ4377 | lJEx1242[plgc-39(2kb)::mKate2::gpd-23'UTR; unc-122::gfp] |
| AQ4428 | lJEx1255[plgc-58(2kb)::mKate2::gpd-23'UTR; unc-122::gfp] |
| AQ5072 | lJEx1562[plgc-46::GFP::gpd-2UTR; punc-122::RFP] |
| OH15262 | "NeuroPAL" otEx7057 |
| AQ4657 | lgc-39(lj121) |
| AQ4850 | NeuroPal(otEx7057); lJEx1453 [plgc-33::lgc-33::SL2 GFP; punc-122::RFP] |
| AQ4837 | NeuroPal(otEx7057); lJEx1449 [plgc-34::lgc-34::SL2 GFP;unc-122::RFP] |
| AQ4529 | him-5(e1490); lJEx1299[plgc-48::gDNA lgc-48 3'UTR::SL2 mKate2;punc-122::GFP] |
| AQ4928 | lJEx1498 [plgc-47(2kb)::lgc-47::SL2 GFP; unc-122::RFP] |
| AQ4849 | NeuroPal(otEx7057); lJEx1453 [plgc-49::lgc-49::SL2 GFP; unc-122::RFP] |

Note:
*lgc-57* previously called *ggr-1*
*lgc-58* previously called *ggr-2*

**Supplementary Table 3:**
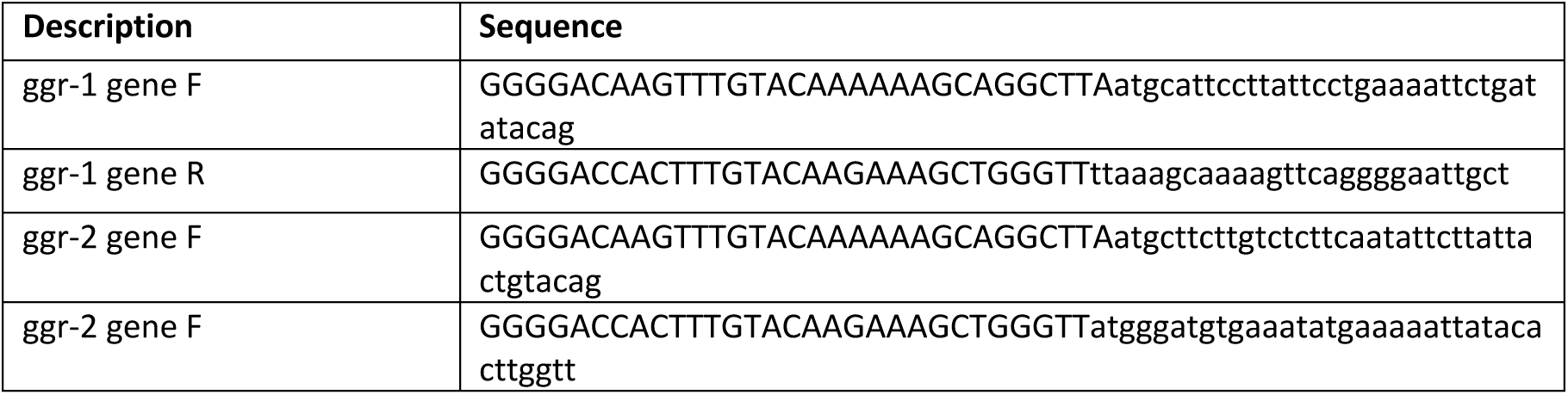
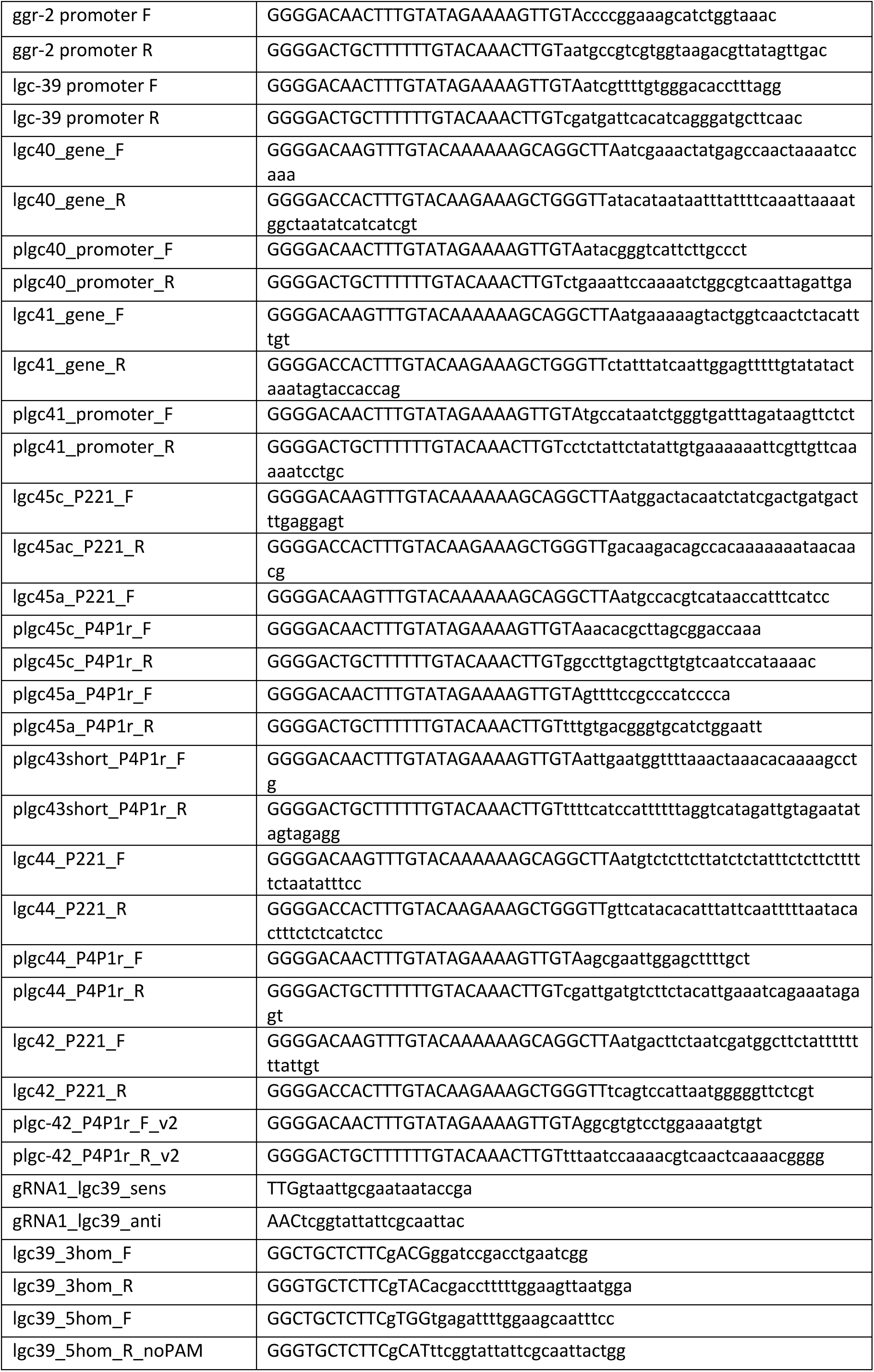
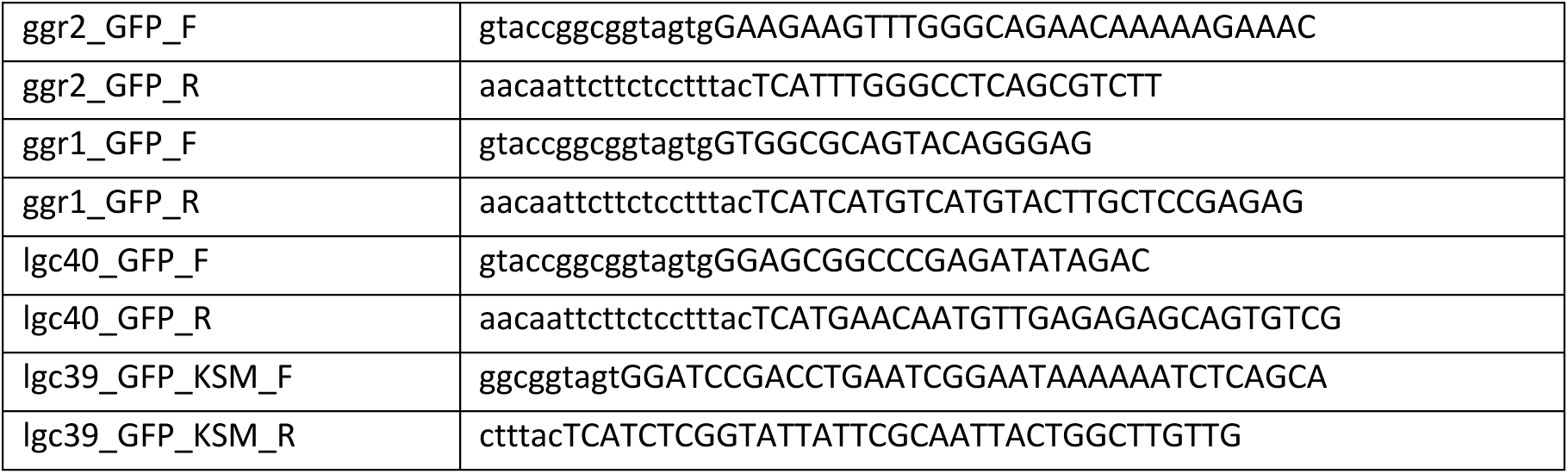
List of primers used in this study.

**Supplementary Figure 1:**
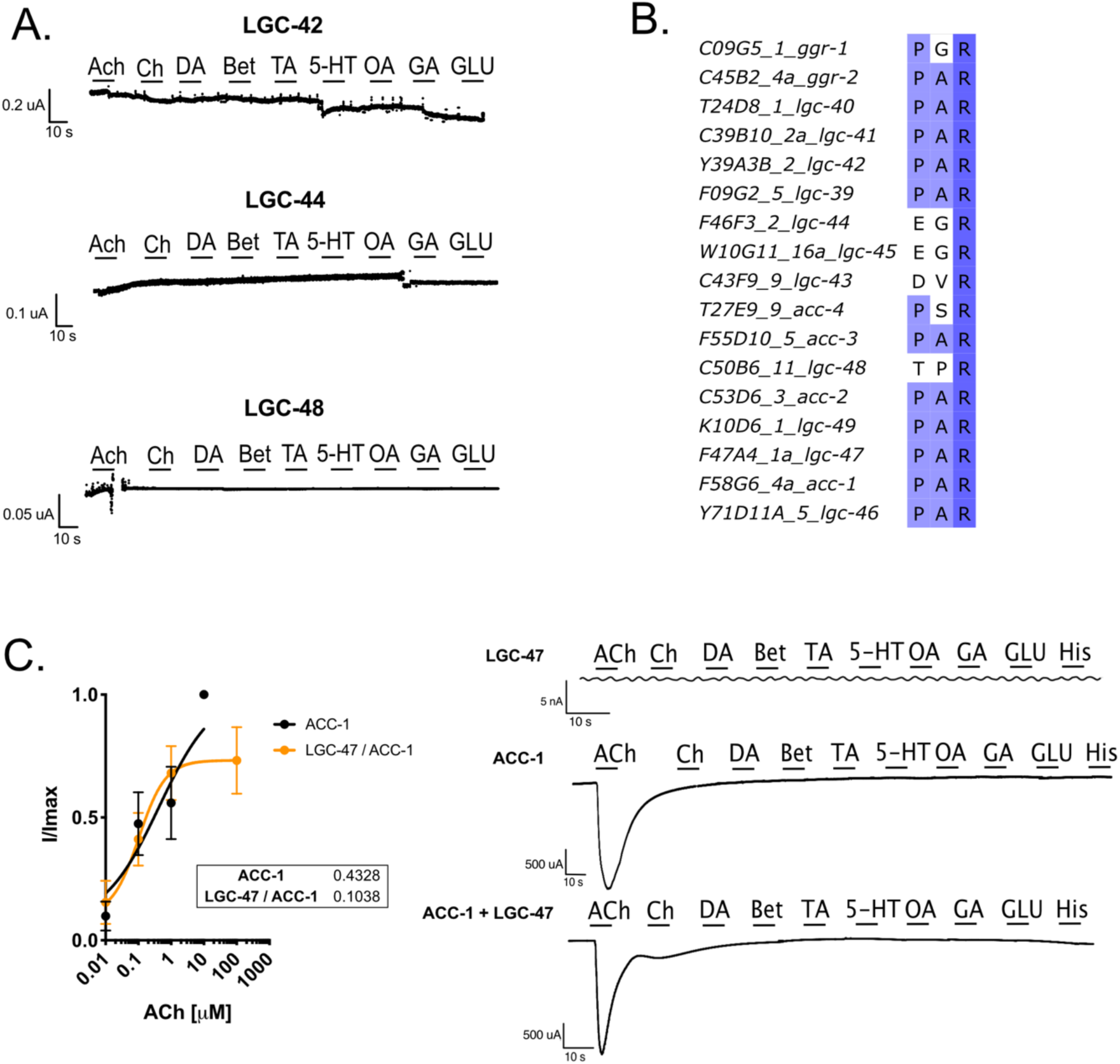
Negative traces for still orphan LGICs, representative ion selectivity curves and channel kinetics. A. Continuous TEVC traces from oocytes clamped at -60mV expressing LGC-42, LGC-44, and LGC-48, exposed to 10s of a panel of ligands. B. Alignment of PAR motifs for channels in the diverse and ACC groups. C. LGC-47 and ACC-1 do not form a heteromeric channel in *Xenopus* oocytes. Left: Acetylcholine-induced dose response curves for oocytes expressing ACC-1 alone, or in combination with LGC-47. Error bars represent SEM of 7-12 oocytes per construct, insert shows EC_50_ values. Right: Continuous TEVC traces from oocytes clamped at -60mV expressing LGC-47, ACC-1, or a combination, exposed to 10s of a panel of ligands. Note that the small changes in current seen in the traces are attributed to recording and perfusion artefacts.

**Supplementary Figure 2:**
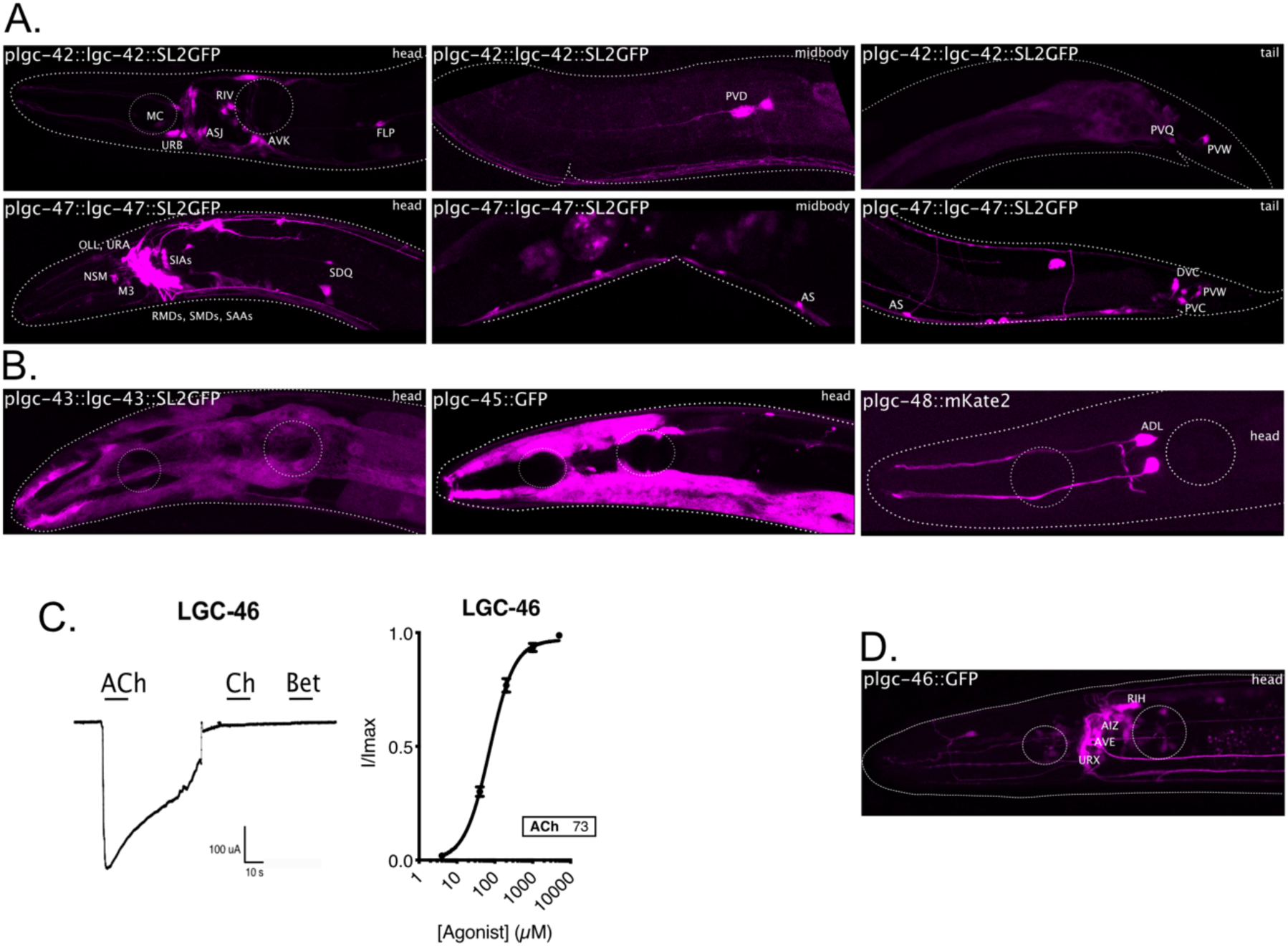
Expression patterns of still orphan LGICs and LGC-46 characterisation. A-B. Expression of fluorescent reporters for still orphan LGCIs, head body and tail are shown for *lgc-42* and *lgc-47* (A), head only is shown for *lgc-43*, *lgc-45* and *lgc-48* (B). C. Acetylcholine-induced dose response curve, continuous trace, and expression pattern of LGC-46. Error bars represent SEM of 6 oocytes. Insert shows EC_50_. Dose response curves for GFP tagged LGICs in response to their major activating ligands. Error bars represent SEM of 4-5 oocytes per construct, inserts show EC_50_. D. Expression of fluorescent reporters for plgc-46 shows a broad neuronal expression pattern with expression in e.g., AIZ, RIH and AVE neurons.

**Supplementary Figure 3:**
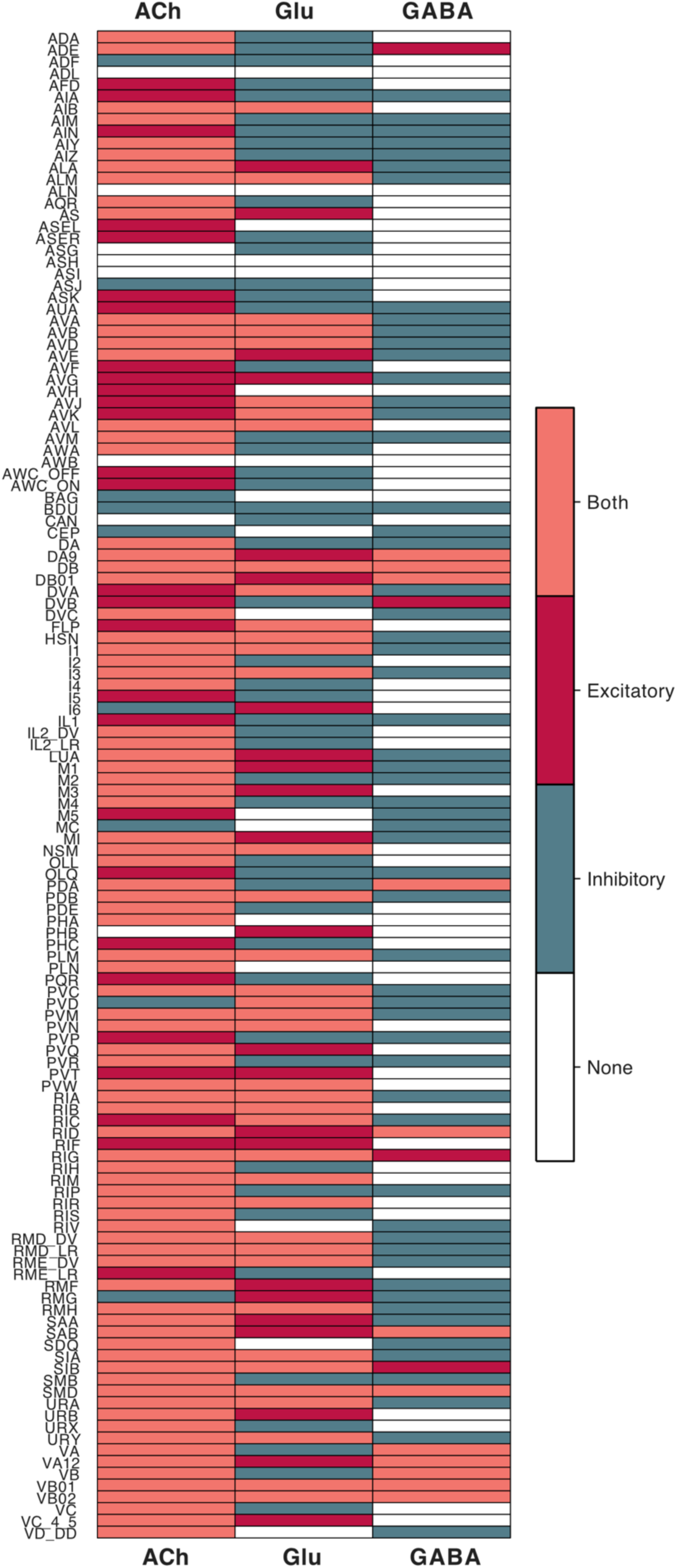
Binary heatmap of synaptic sign prediction for the three major neurotransmitters; ACh, glutamate and GABA. The heatmap shows the summed expression level of all LGICs in *C. elegans* per neural class and neurotransmitter, a net sum of excitatory channels is displayed in red, inhibitory in green, equal expression in peach and no expression of LGICs in white.

**Supplementary Figure 4:**
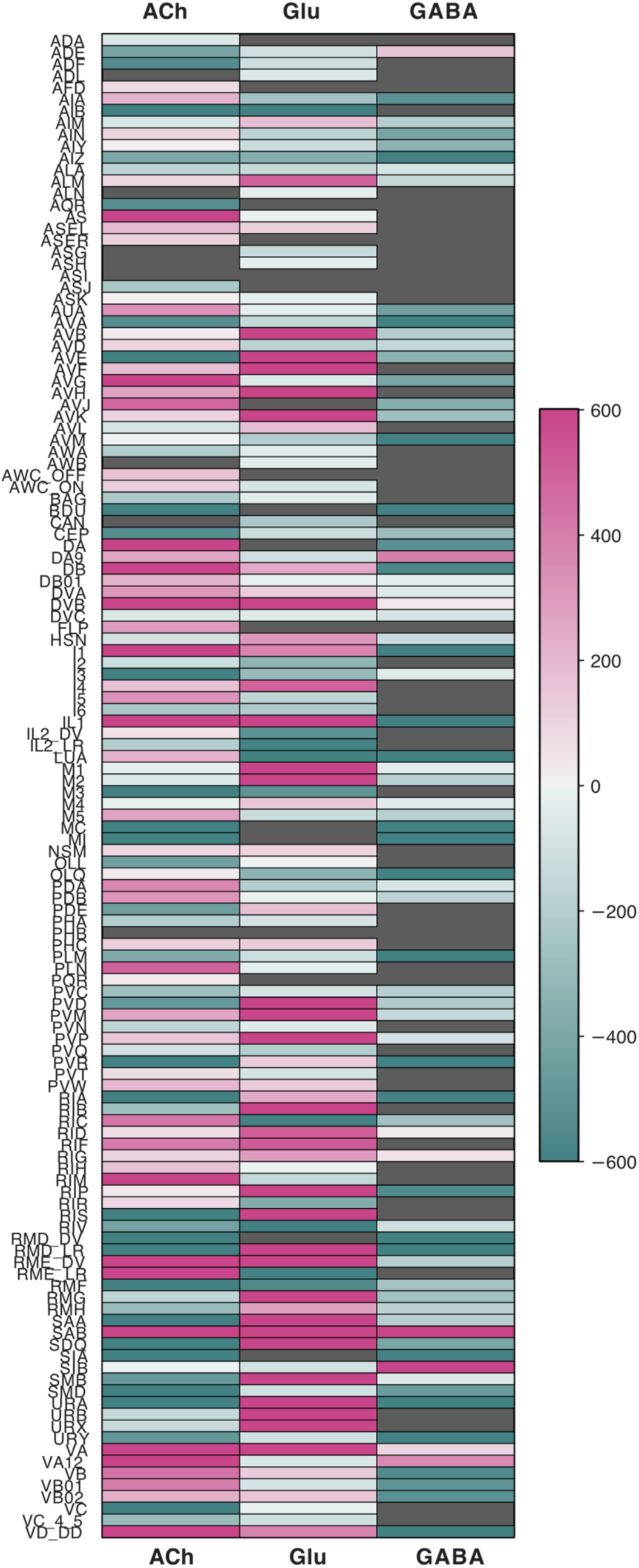
Expression heatmap for the three major neurotransmitters. The heatmap shows the summed expression level, and ion selectivity, for LGICs separated by transmitter (ACh, glutamate and GABA) and neural class. Net excitatory channel expression is represented in pink and inhibitory in green.

**Supplementary Figure 5:**
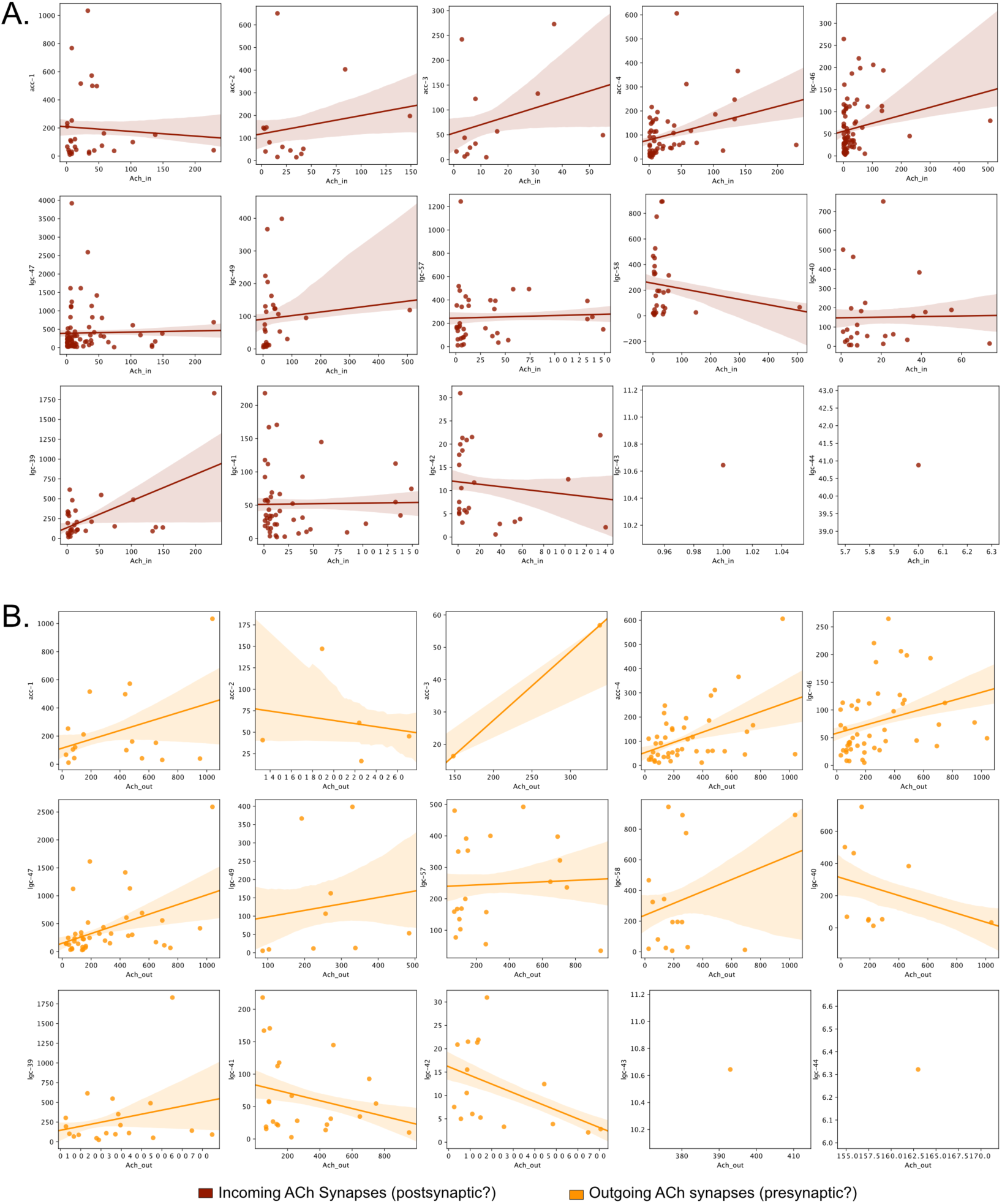
Correlation graphs between cholinergic synapses and expression of selected LGICs. Scatter plots showing gene expression level vs total number of incoming cholinergic synapses (red) and outgoing cholinergic synapses (orange). Lines fit using relplot with shaded areas representing the standard error of the line fit.

**Supplementary Figure 6:**
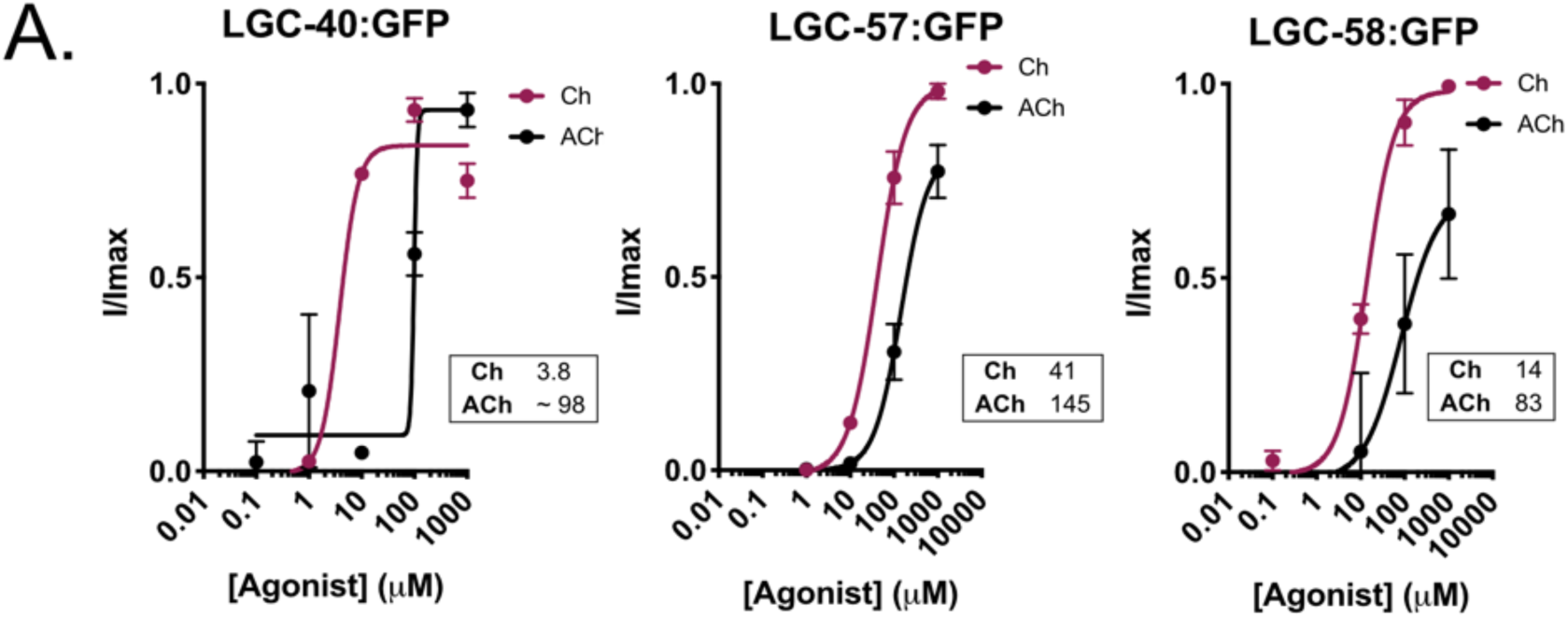
GFP tagging of LGICs does not influence channel function. Acetylcholine- or choline-induced dose response curves for oocytes expressing GFP tagged versions of LGC-40, LGC-57 and LGC-58 shows that EC_50_ values are not influenced by the insertion of the GFP tag. Error bars represent SEM of 7-12 oocytes per construct, insert shows EC_50_ values.

